# Proteasome Inhibition Enhances Lysosome-mediated Targeted Protein Degradation

**DOI:** 10.1101/2025.01.31.634950

**Authors:** Ahmed M. Elshazly, Nayyerehalsadat Hosseini, Janakiram Vangala, Shanwei Shen, Victoria Neely, Xiaoyan Hu, Piyusha P. Pagare, Hisashi Harada, Steven Grant, Senthil K. Radhakrishnan

## Abstract

Proteasome inhibitor drugs are currently used in the clinic to treat multiple myeloma and mantle cell lymphoma. These inhibitors cause accumulation of undegraded proteins, thus inducing proteotoxic stress and consequent cell death. However, cancer cells counteract this effect by activating an adaptive response through the transcription factor Nuclear factor erythroid 2-related factor 1 (NRF1, also known as NFE2L1). NRF1 induces transcriptional upregulation of proteasome and autophagy/lysosomal genes, thereby reducing proteotoxic stress and diminishing the effectiveness of proteasome inhibition. While suppressing this protective autophagy response is one potential strategy, here we investigated whether this heightened autophagy could instead be leveraged therapeutically. To this end, we designed an autophagy-targeting chimera (AUTAC) compound to selectively degrade the anti-apoptotic protein Mcl1 via the lysosome. Our results show that this lysosome-mediated targeted degradation is significantly amplified in the presence of proteasome inhibition, in a NRF1-dependent manner. Mechanistically, AUTAC-driven Mcl1 clearance requires K63-linked ubiquitination by UBC13 and TRAF6 and recognition by the cargo receptor p62/SQSTM1. The combination of the proteasome inhibitor carfilzomib and Mcl1 AUTAC synergistically promoted cell death in both in vitro models, including wild-type and proteasome inhibitor-resistant multiple myeloma and lung cancer cells, and in mouse tumor xenografts. Thus, our work offers a novel strategy for enhancing proteasome inhibitor efficacy by exploiting the adaptive autophagy response. More broadly, our study establishes a framework for amplifying lysosome-mediated targeted protein degradation, with potential applications in cancer therapeutics and beyond.

## INTRODUCTION

Proteasome inhibitors (PIs) are a cornerstone in the treatment of multiple myeloma and are also part of mantle cell lymphoma therapeutic protocols ^1,2^. FDA-approved PIs bortezomib, carfilzomib, and ixazomib are small-molecule inhibitors designed to target the proteasome. Bortezomib and ixazomib are peptide boronate inhibitors, mimicking proteasome substrates through dipeptide-like structures, while carfilzomib is an epoxyketone-based inhibitor that binds irreversibly to the β5 subunit ^3^. By blocking proteasomal activity, these drugs induce endoplasmic reticulum (ER) stress due to the accumulation of unfolded proteins and insoluble aggregates, which ultimately leads to cell cycle arrest, activation of antiproliferative signals, and apoptosis ^1,4,5^. While PIs have demonstrated clinical success in blood malignancies, they have shown limited efficacy against solid tumors ^6–11^. Additionally, their long-term utility is often hindered by the development of resistance, leading to relapse and poor prognosis ^12^.

Tumor cells activate a range of adaptive mechanisms to counteract the cytotoxic effects of PIs, including the activation of nuclear factor erythroid 2-related factor 1 (NRF1, also known as NFE2L1). NRF1 resides in the ER membrane as an inactive p120 precursor and is cleaved by the protease DDI2 to its active p110 form upon proteasome inhibition. The active NRF1 translocates to the nucleus, where it upregulates the expression of proteasome and autophagy/lysosomal genes, thereby mitigating proteotoxic stress and reducing PI efficacy ^13–15^. Although directly targeting NRF1 remains challenging due to the complexity of its pathway, adaptive responses such as NRF1-mediated autophagy induction could represent potential therapeutic vulnerabilities. Autophagy, a cytoplasmic recycling process critical for maintaining cellular homeostasis, enables tumor cells to survive under chemotherapeutic stress - a phenomenon referred to as cytoprotective autophagy ^16,17^. Multiple studies have linked autophagy activation to PI resistance development ^18–21^. Instead of inhibiting autophagy, here we sought to exploit this cytoprotective mechanism for therapeutic benefit using autophagy-targeting chimera (AUTAC) technology. AUTACs rely on cGMP-mediated S-guanylation, which has been implicated in the selective autophagic degradation of Group A Streptococcus (GAS) ^22^. These compounds consist of an autophagy degradation tag, typically fluorobenzyl guanine (FBnG) derivatives, linked to a ligand that targets a specific protein of interest ^23^.

The anti-apoptotic protein Mcl1, a key regulator of mitochondrial integrity, binds and sequesters pro-apoptotic proteins such as Bax, Bak, and Bok, thereby preventing cytochrome C release and apoptosis ^24,25^. Elevated Mcl1 levels following proteasome inhibition contribute to PI resistance in multiple myeloma ^26–28^. Preclinical studies have shown that suppressing Mcl1 restores PI sensitivity ^27,29–32^, and several clinical trials are underway to evaluate Mcl1 inhibitors in multiple myeloma (NCT04178902, NCT03465540, NCT04543305, NCT02675452, NCT02992483). However, these efforts are constrained by concerns over potential cardiac toxicity ^33^.

In this study, we developed an AUTAC compound to selectively degrade Mcl1 via the lysosomal pathway. Our results demonstrate that this lysosome-mediated targeted degradation is significantly enhanced by proteasome inhibition in an NRF1-dependent manner. The combination of carfilzomib and Mcl1 AUTAC synergistically induced cell death in both wild-type and PI-resistant multiple myeloma and non-small cell lung cancer cells. Importantly, AUTAC-mediated potentiation of carfilzomib cytotoxicity was further validated in a mouse xenograft tumor model, where the combination treatment resulted in significantly enhanced antitumor efficacy compared with either agent alone. Overall, our findings provide a novel strategy to enhance PI efficacy by leveraging the adaptive autophagy response.

## RESULTS

### AUTAC-Mediated Degradation of Mcl1 via Autophagic Machinery

Building on the efforts of Arimoto’s group ^22,23,34^ and recognizing the importance of targeting Mcl1, we developed an Mcl1-targeting AUTAC utilizing a second-generation pyrazole-linked FBnG tag. This tag was conjugated via a pyrazole linker to the S1-6 derivative, a well-characterized Mcl1 inhibitor ^35^ (Fig. 1A). Sequence alignment revealed moderate overall conservation between Mcl1 and Bcl2 (33% identity, 50% similarity) and between Mcl1 and Bclxl (29% identity, 44% similarity), with notable divergence localized near the periphery of the BH3-binding groove (Supplementary Fig. 1A and B). These differences suggested that local surface topology at the groove entrance might influence accommodation of the AUTAC linker and degradation tag. To investigate this possibility, molecular docking studies were performed using Schrödinger’s Glide module ^36,37^. Crystal structures of Mcl1 (PDB ID 6YBL^38^), Bcl2 (PDB ID 2XA0 ^39^), and Bclxl (PDB ID 1R2D ^40^) were prepared using standard protocols, and the isolated warhead was first docked to establish a common binding framework. In all three proteins, the warhead adopted a conserved pose within the canonical BH3-binding groove (Supplementary Fig. 1C), consistent with the known BH3-mimetic ligand binding modes ^41,42^. Importantly, the orientation of the linker exit position was preserved across targets, indicating that any differences in degradation are unlikely to arise from impaired warhead engagement.

**Figure 1.**
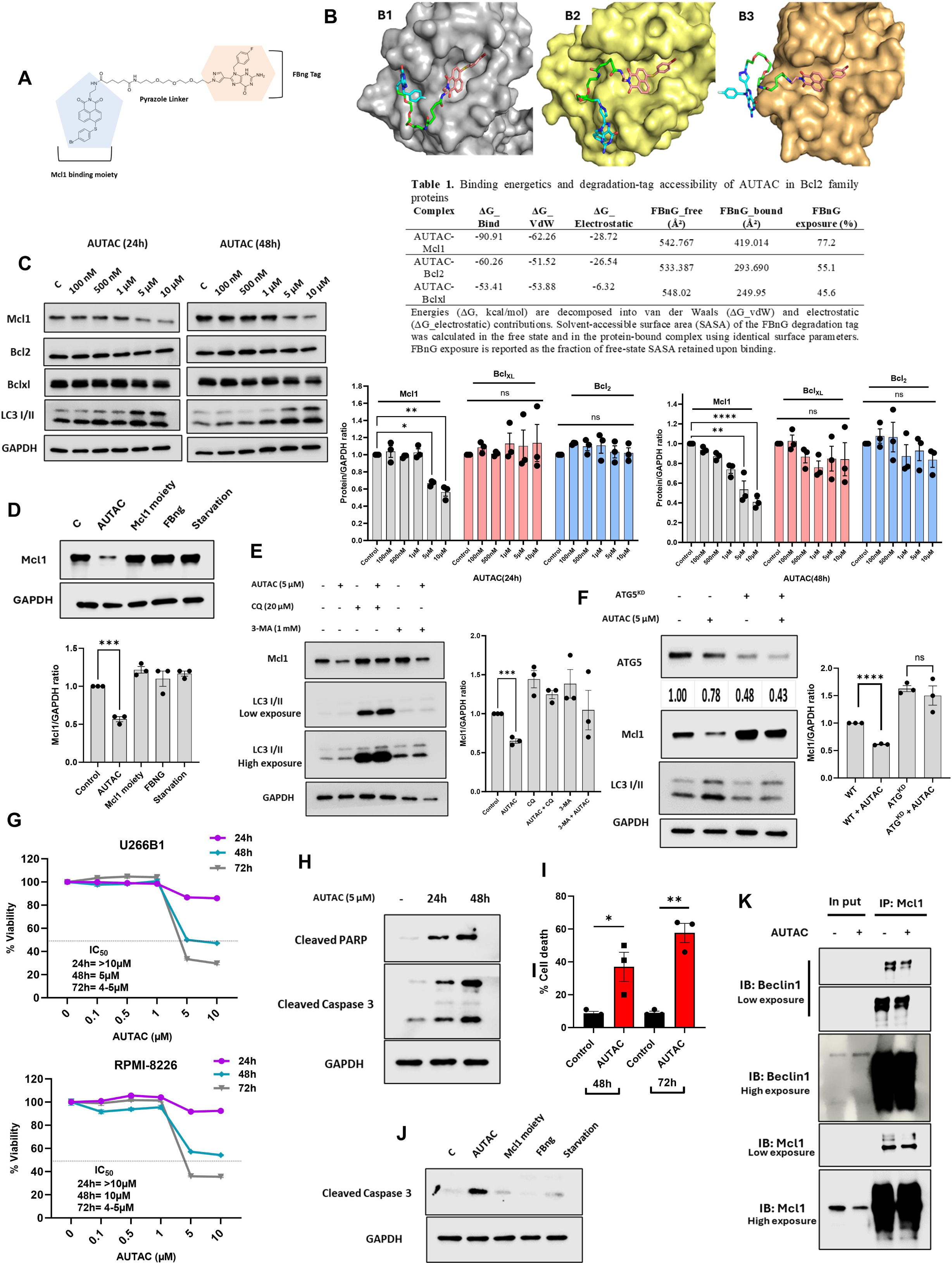
AUTAC degrades Mcl1 through autophagy. **(A)** The chemical structure of Mcl1-specific AUTAC consists of a Mcl1 binding moiety connected to a FBnG-tag via a pyrazole linker. The second-generation AUTAC achieved superior degradative efficiency by overcoming the membrane permeability limitations associated with the negatively charged cyclic phosphate scaffold of the first-generation construct. **(B)** AUTAC binding in (B1) Mcl1, (B2) Bcl2 and (**3**) Bclxl. Mcl1 (PDB ID: 6YBL), Bcl2 (PDB ID: 2XA0) and Bclxl (PDB ID: 1R2D) are shown as grey, yellow and orange surfaces. Warhead, linker and FBnG tag are shown as salmon, green and cyan sticks respectively. **(C)** Western blot analysis for Mcl1, Bcl2, Bclxl, and LC3I/II proteins in U266B1 cells treated with the indicated concentrations of AUTAC for 24h and 48h. GAPDH was used as the loading control. **(D)** Western blot analysis for Mcl1 protein in U266B1 cells treated with AUTAC (5 μM), Mcl1 moiety (5 μM), FBnG (10 μM), as well as starved U266B1 cells for 24h. GAPDH was used as the loading control. **(E)** Western blot analysis for Mcl1, and LC3I/II proteins in U266B1 cells treated with different combinations of AUTAC (5 μM), CQ (20 μM) or 3-MA (1 mM) for 24h. GAPDH was used as the loading control. **(F)** Western blot analysis for ATG5, Mcl1, and LC3I/II proteins in U266B1 wild-type and U266B1 ATG5^KD^ cells treated with AUTAC (5 μM) for 24h. GAPDH was used as the loading control. **(G)**The percentage of cellular viability for U266B1 and RPMI-8226 cells treated with the indicated concentrations AUTAC for 24, 48 and 72h. **(H)** Western blot analysis for cleaved caspase 3 and cleaved PARP proteins in U266B1 cells treated with AUTAC (5 μM) for 24 and 48h. GAPDH was used as the loading control. **(I)** The percentage of cell death for U266B1 cells treated with AUTAC (5 μM) for 48 and 72h. **(J)** Western blot analysis for cleaved caspase 3 protein in U266B1 cells treated with AUTAC (5 μM), Mcl1 moiety (5 μM), FBnG (5 μM), as well as starved U266B1 cells for 24h. GAPDH was used as the loading control. **(K)** Western blot analysis of Mcl1 and Beclin1 in U266B1 cells treated with AUTAC (10 μM) for 24 h, followed by co-immunoprecipitation. Three biological replicates for each cell line were used to perform cell viability assay and Western blotting. The uncropped blots are shown in the Supplemental Materials. Statistical significance of each condition compared to the indicated control or treatment was determined using unpaired Student’s t-test or two-way ANOVA with Tukey or Sidak post hoc tests, as appropriate. Data are represented as mean ± SEM. Significance levels are indicated as follows: * p < 0.05, ** p < 0.01, *** p < 0.001 and **** p < 0.0001.

AUTAC docking was then carried out using a core-constrained protocol to maintain the validated warhead pose while allowing flexibility of the linker and FBnG degradation tag. Across all three proteins, clustering analysis revealed a dominant binding family that differed primarily in linker trajectory and FBnG positioning at the groove periphery (Fig. 1B). Docking scores favored Mcl1 (top GlideScore −10.74) over Bcl2 (−7.10) and Bclxl (−7.13), despite identical warhead constraints, suggesting that selectivity arises from differential accommodation of the bifunctional architecture rather than intrinsic warhead affinity. Structural inspection revealed clear topological differences at the groove entrance. In the Mcl1 complex, the PEG linker extended along a shallow, solvent-exposed surface channel, positioning the FBnG tag away from the protein surface and largely accessible to solvent (Fig. 1B). In contrast, both Bcl2 and Bclxl presented more concave and restrictive surface contours, causing the linker to pack tightly against the protein and directing the FBnG tag into recessed surface regions near the groove entrance (Fig. 1B). These geometries suggest that Mcl1 uniquely supports productive presentation of the degradation tag.

To further quantify these differences, the top five docking poses for each target were subjected to Prime MM-GBSA^43^ rescoring. AUTAC binding to Mcl1 was energetically favored (average ΔG_bind ≈ −90.9 kcal/mol) relative to Bcl2 (−60.3 kcal/mol) and Bclxl (−53.4 kcal/mol) (Table 1). This preference was driven primarily by more favorable van der Waals interactions in Mcl1, with additional contributions from electrostatic complementarity, which was markedly reduced in Bclxl. Because productive AUTAC function requires solvent accessibility of the degradation tag, solvent-accessible surface area (SASA) analysis ^44^ was performed on the FBnG moiety. In the highest-ranked Mcl1 complex, FBnG retained 77% of its free-state SASA, whereas exposure was substantially reduced in Bcl2 (55%) and Bclxl (46%) (Table 1). Thus, despite preserved warhead binding, FBnG becomes progressively more buried in Bcl2- and Bclxl -bound conformations. Together, these results support a model in which selective Mcl1 targeting arises from favorable surface topology at the BH3-groove periphery that permits solvent-exposed presentation of the degradation tag. In contrast, steric confinement in Bcl2 and Bclxl limits FBnG accessibility, providing a structural rationale for preferential Mcl1 targeting despite conserved warhead engagement.

To further verify the selectivity of AUTAC, we evaluated its effect on Bcl2 family proteins in the U266B1 cell line across various concentrations (Fig. 1C). AUTAC reduced Mcl1 protein levels in a dose- and time-dependent manner without an observable impact on both Bclxl and Bcl2. Additionally, AUTAC increased LC3 I/II levels, indicating activation of the autophagic machinery (Fig. 1C). To assess whether AUTAC induces non-selective autophagy through which Mcl1 may be degraded, we compared its effects with starvation-induced autophagy, an approach known to trigger non-selective protein degradation ^45–47^ (Supplementary Fig. 1D and E). Initially, we validated that autophagy was induced by starvation using LysoTracker staining (Supplementary Fig. 1D). To further confirm autophagy induction, we performed an autophagic flux assay (Supplementary Fig. 1E). Treatment with chloroquine (CQ) in combination with starvation led to greater accumulation of LC3II compared to CQ alone, indicating an active autophagic activity. Under starvation, Mcl1 levels remained largely unchanged (Fig. 1D), suggesting that Mcl1 is not efficiently targeted by non-selective autophagy. Importantly, the individual components of AUTAC (FBng and the Mcl1-binding moiety) did not affect Mcl1 expression when applied separately (Fig. 1D), confirming that the degradation of Mcl1 is dependent on the full AUTAC construct and indicates its selectivity. We further evaluated the selectivity of the AUTAC for Mcl1 degradation by examining several unrelated proteins, including mTOR, p53, HSP70, HSP60, ERK, and ULK1, whose protein levels were not affected by AUTAC treatment (Supplementary Fig. 2A). These results were further supported by immunofluorescence staining of U266B1 cells treated with AUTAC for 24 and 48 hours, which revealed a time-dependent reduction in Mcl1 levels, accompanied by LC3B and LAMP1 accumulation. Notably, increased colocalization between Mcl1 and LAMP1 or LC3B was observed at 24 hours (Supplementary Fig. 2B).

To confirm the autophagy-dependent mechanism of AUTAC, we pharmacologically inhibited the autophagic machinery using CQ ^48^ and 3-Methyladenine (3-MA) ^49^. This inhibition rescued AUTAC-mediated degradation of Mcl1 in U266B1 cells (Fig. 1E). Moreover, combining AUTAC with CQ led to greater LC3II accumulation compared to CQ alone, consistent with enhanced autophagic flux (Fig. 1E). Genetic knockdown of ATG5 similarly alleviated AUTAC-mediated Mcl1 degradation (Fig. 1F). These findings establish that AUTAC selectively targets Mcl1 for degradation through autophagy.

We next examined the functional impact of AUTAC-mediated Mcl1 degradation on cell viability. Treatment with AUTAC reduced the viability of U266B1 and RPMI-8226 cells in a time-dependent manner with an IC of approximately 5µM after 48 h (Fig. 1G). To determine whether this reduction resulted from inhibited proliferation or induction of apoptosis, we performed western blot analysis for cleaved caspase 3 and cleaved PARP. AUTAC treatment led to robust accumulation of both cleaved caspase 3 and cleaved PARP at 24 and 48 hours (Fig. 2H), indicating activation of apoptotic pathways. These findings were corroborated by a trypan blue exclusion assay, which showed that AUTAC induced approximately 50% cell death at 48 hours, increasing to around 60% by 72 hours (Fig. 2I). Collectively, these data demonstrate that AUTAC induces apoptotic cell death in U266B1 cells. To confirm the specificity of this effect, we assessed whether the individual components of AUTAC (FBnG or the Mcl1-binding moiety) were capable of inducing apoptosis on their own. Neither component, when applied separately, nor 24-hour starvation treatment resulted in caspase 3 cleavage (Fig. 2J). These findings indicate that both Mcl1 binding and the autophagy-targeting function are required for AUTAC’s activity, highlighting the selectivity and functional integrity of the full AUTAC construct.

**Figure 2.**
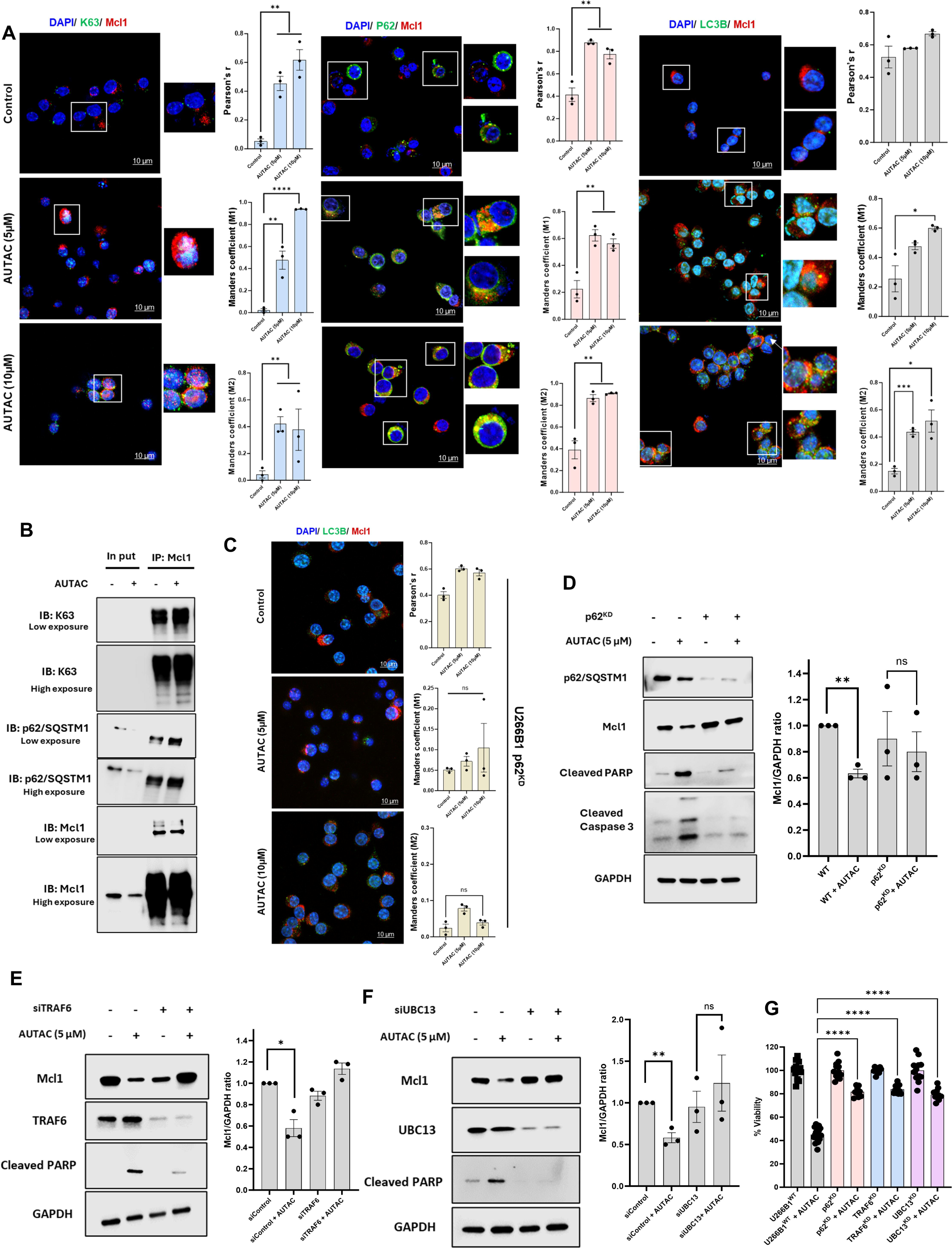
AUTAC selectively degrades Mcl1 via TRAF6/UBC13/p62(SQSTMI). **(A)** Confocal images for Mcl1, K63, p62/SQSTM1 and LC3B proteins in U266B1^WT^ cells treated with AUTAC (5 and 10 μM) for 24 (scale 10 μM). **(B)** Western blot analysis of Mcl1, K63, and p62/SQSTM1 in U266B1 cells treated with AUTAC (10 μM) for 16 h, followed by co-immunoprecipitation. **(C)** Confocal images for Mcl1, and LC3B proteins in U266B1p62^KD^ cells treated with AUTAC (5 and 10 μM) for 24 (scale 10 μM). **(D)** Western blot analysis for p62/SQSTM1, Mcl1 and cleaved PARP and cleaved caspase 3 proteins in U266B1^WT^ and U266B1 p62^KD^ cells treated with AUTAC (5 μM) for 24h. GAPDH was used as the loading control. **(E and F)** Western blot analysis for Mcl1, TRAF6, UBC13 and cleaved PARP proteins in U266B1^WT^, U266B1 TRAF6 ^KD^, and UBC13 ^KD^ treated with AUTAC (5 μM) for 24h. GAPDH was used as the loading control. **(G)** The percentage of cell death for U266B1^WT^, p62^KD^, TRAF6^KD^, and UBC13^KD^ cells treated with AUTAC (5 μM) for 72h. Three biological replicates for each cell line were used to perform cell viability assay and Western blotting. The uncropped blots are shown in the Supplemental Materials. Statistical significance of each condition compared to the indicated control or treatment was determined using unpaired Student’s t-test or two-way ANOVA with Tukey or Sidak post hoc tests, as appropriate. Data are represented as mean ± SEM. Significance levels are indicated as follows: * p < 0.05, ** p < 0.01, *** p < 0.001 and **** p < 0.0001.

To elucidate the mechanism underlying AUTAC-induced autophagy, we first excluded the possibility that Mcl1 AUTAC generally induce autophagy by comparing the Mcl1 AUTAC with an unrelated MetAP2-targeting AUTAC. The MetAP2 AUTAC did not affect Mcl1 levels and failed to induce autophagy, in contrast to the Mcl1 AUTAC (Supplementary Fig. 3A). These findings were further confirmed by autophagic flux inhibition assays (Supplementary Fig. 3B). We next excluded the involvement of the mTOR pathway by comparing the effects of Mcl1 AUTAC with rapamycin. While rapamycin clearly inhibited phosphorylated mTOR (p-mTOR) and the phosphorylation of its downstream target S6 and induced autophagy, Mcl1 AUTAC did not affect the phosphorylation status of these targets (Supplementary Fig. 3C and D). Furthermore, rapamycin did not alter Mcl1 protein levels, whereas AUTAC led to a significant reduction in Mcl1 protein level (Supplementary Fig. 3C). We further showed that the intact AUTAC molecule, but not its individual components—the Mcl1-binding moiety or FBnG—induces autophagy (Supplementary Fig. 3E), confirming that AUTAC-mediated autophagy is a selective process rather than a nonspecific induction of general autophagy. Based on these findings, we hypothesized that AUTAC promotes autophagy via targeted degradation of Mcl1. First, we demonstrated that AUTAC-mediated autophagy is dependent on Mcl1. Knockdown of Mcl1 markedly abrogated AUTAC-induced autophagic activity, as evidenced by diminished LC3II accumulation following co-treatment with AUTAC and CQ in Mcl1^KD^ cells compared with their control counterparts (Supplementary Fig. 3F). Two main molecular mechanisms have been discussed in the scientific literature: (1) Mcl1 is essential for maintaining mitochondrial integrity, and its inhibition leads to increased reactive oxygen species (ROS) ^50,51^, which in turn induce autophagy ^52,53^; and (2) degradation of Mcl1 leads to dissociation from Beclin1, thereby activating Beclin1-dependent autophagy ^54,55^. To test the first hypothesis, we treated cells with AUTAC in the presence of N-acetylcysteine (NAC), a known ROS scavenger^56^, and assessed autophagic flux. While AUTAC in combination with CQ led to greater accumulation of LC3II compared to CQ alone, NAC treatment did not block this effect (Supplementary Fig. 3G). This suggests that AUTAC-mediated autophagic flux is not dependent on ROS generation. To investigate the second hypothesis, we performed co-immunoprecipitation to assess whether AUTAC affects the interaction between Mcl1 and Beclin1 (Fig. 1K). Interestingly, AUTAC reduced the binding between Mcl1 and Beclin1 (Fig. 1K) and increased the phosphorylation of Beclin1 (Supplementary Fig. 3H), suggesting that AUTAC liberates Beclin1, leading to its activation and subsequent induction of autophagy.

Collectively, we provide comprehensive evidence demonstrating that AUTAC selectively targets Mcl1, induces cell death in multiple myeloma cell lines, and triggers a specific, Beclin1-dependent autophagy process.

### AUTAC selectively degrades Mcl1 through TRAF6/UBC13/p62(SQSTM1) axis

Takahashi et al. ^23^ demonstrated that AUTAC technology promotes K63-linked polyubiquitination of target proteins. The selective autophagy receptor, p62/SQSTM1, recognizes and binds these K63-linked ubiquitin chains via its UBA (Ubiquitin-Associated) domain ^57,58^. Through its LIR (LC3-interacting region) motif, p62 subsequently interacts with LC3 on the autophagosome membrane ^59,60^, facilitating the selective degradation of ubiquitinated cargo by the autophagic machinery. To investigate whether our AUTAC compound operates through this mechanism, we performed immunofluorescence staining on U266B1 cells treated with AUTAC at 5 and 10 μM for 24 hours. We observed clear colocalization of Mcl1 with K63, p62, and LC3B following AUTAC treatment (Fig. 2A). We further confirmed these results by co-immunoprecipitation, which showed that AUTAC increased the binding of Mcl1 to both K63-linked ubiquitin and p62/SQSTM1 (Fig. 2B). Notably, knockdown of p62/SQSTM1 disrupted the colocalization between LC3B and Mcl1 (Fig. 2C), suggesting that AUTAC mediates K63-linked ubiquitination of Mcl1, which is then recognized and delivered to LC3 by p62 for selective autophagic degradation. To further confirm the p62/SQSTM1-dependent mechanism of action, U266B1^WT^ and U266B1 p62^KD^ cells treated with AUTAC (5 μM) for 24h, where AUTAC-mediated degradation of Mcl1 was abolished upon p62 knockdown compared to its parental counterpart (Fig. 2D). Furthermore, knockdown of p62 abrogated AUTAC-induced accumulation of cleaved caspase 3 and cleaved PARP, indicating that p62 is required for AUTAC-mediated apoptotic signaling (Fig. 2D). These results strongly support the conclusion that AUTAC induces selective autophagic degradation of Mcl1 via the cargo receptor p62/SQSTM1.

Then, we investigated which ligases are involved in AUTAC-mediated Mcl1 degradation. Choi et al. ^61^ previously demonstrated that Mcl1 undergoes K63-linked ubiquitination mediated by TRAF6. Building on this, we postulated that TRAF6, in concert with the E2 conjugating enzyme UBC13, orchestrates AUTAC-induced K63 ubiquitination of Mcl1. Pharmacological inhibition with either C25-140, a selective TRAF6-UBC13 interaction disruptor ^62^, or NSC697923, a potent UBC13 inhibitor ^63^, completely abrogated AUTAC-driven Mcl1 degradation (Supplementary Fig. 4A). Complementary genetic approaches corroborated these findings, as siRNA-mediated depletion of TRAF6 or UBC13 markedly suppressed AUTAC-induced Mcl1 degradation and significantly diminished apoptosis, as evidenced by reduced PARP cleavage (Fig. 2E and F). Together, these results establish TRAF6 and UBC13 as essential mediators of AUTAC-driven K63 ubiquitination and degradation of Mcl1, thereby underscoring the mechanistic selectivity of AUTAC in targeting this anti-apoptotic protein.

To further delineate the mechanism underlying AUTAC-induced cytotoxicity, we assessed whether selective degradation is required for its anti-tumor activity. AUTAC treatment alone reduced U266B1^WT^ cell viability by more than 50%. However, knockdown of p62, TRAF6, or UBC13 substantially blunted this cytotoxic response (Fig. 2G). These results demonstrate that AUTAC-driven cell death is mechanistically dependent on targeted Mcl1 degradation and strongly support the selectivity and pathway fidelity of the AUTAC system.

### Autophagy Induction Augments the AUTAC-Mediated Degradation of Mcl1

We hypothesized that combining an autophagy inducer with AUTAC could enhance its ability to degrade Mcl1. To test this notion, we induced autophagy in U266B1 cells via starvation ^64^, which was confirmed by lysotracker staining, the autophagic flux inhibition assay and the degradation of p62 (Supplementary Fig. 1D, E and 4B). Treatment of U266B1 cells with 5 μM AUTAC for 24 hours in conjunction with starvation resulted in more pronounced Mcl1 degradation compared to AUTAC alone (Supplementary Fig. 4B).

Carfilzomib (CFZ), a first-line proteasome inhibitor for relapsed and refractory multiple myeloma ^65^, has been shown by our group and others to induce autophagy ^13,18,19,66^. Consistent with these reports, CFZ treatment induced autophagy in U266B1, MM.1S, RPMI-8226 cells, as evidenced by lysotracker staining and the autophagic flux inhibition assay, (Supplementary Fig. 5A, B and C). This induction of autophagic flux was also observed in the proteasome inhibitor-resistant U266B1^R^ cells (Supplementary Fig. 5C). We next evaluated the combined effect of CFZ and AUTAC on Mcl1 levels. CFZ enhanced AUTAC-mediated Mcl1 degradation across multiple myeloma cell lines, including U266B1, MM.1S, RPMI-8226, and AMO-1, with a concurrent increase in autophagic flux, as shown by LC3I/II levels (Fig. 3A). Furthermore, this effect was recapitulated in proteasome inhibitor-resistant multiple myeloma cell lines, such as U266B1^R^ and AMO-1^R^ (Fig. 3B). Notably, LC3II levels differed significantly ^67^, particularly in MM.1S and RPMI-8226 cells. To determine whether this reflected ongoing autophagic activity, we conducted autophagic flux inhibition assay in the presence of CQ. CQ in combination with CFZ led to a marked increase in LC3II levels in both cell lines as compared to CQ alone (Supplementary Fig. 5B), indicating that LC3II is actively degraded ^67^. These results confirm the presence of a highly dynamic and functional autophagic flux.

**Figure 3.**
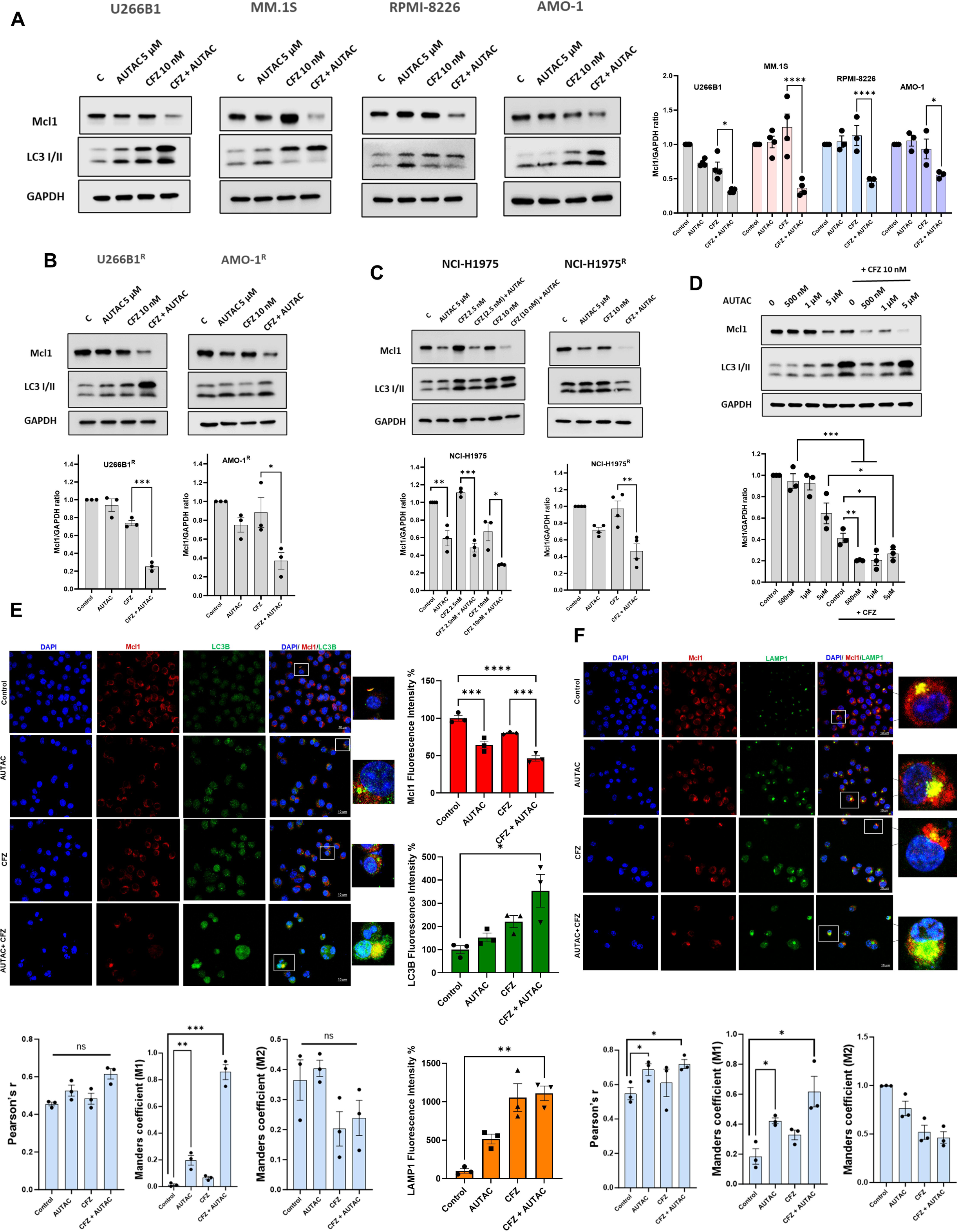
Carfilzomib enhances AUTAC-mediated Mcl1 degradation in multiple myeloma and non-small cell lung cancer cells. **(A)** Western blot analysis for Mcl1, and LC3I/II proteins in U266B1, MM.1S, RPMI-8226 and AMO-1 cells treated with AUTAC (5 μM) and/or CFZ (10 nM) for 16h, except for AMO-1 cells treated for 24h. GAPDH was used as the loading control. **(B)** Western blot analysis for Mcl1, and LC3I/II proteins in U266B1^R^ and AMO-1^R^ cells treated with AUTAC (5 μM) for 48h, then CFZ (10 nM) was added for the last 24h. GAPDH was used as the loading control. **(C)** Western blot analysis for Mcl1, and LC3I/II proteins in NCI-H1975 and NCI-H1975^R^ cells treated with AUTAC (5 μM) and/or CFZ (2.5 nM, and 10 nM) for 72h. GAPDH was used as the loading control. **(D)** Western blot analysis for Mcl1, and LC3I/II proteins in U266B1 cells treated with the indicated concentrations of AUTAC and/or CFZ (10 nM) for 16h. GAPDH was used as the loading control. **(E and F)** Confocal images and their quantification for Mcl1, LC3B **(E)** and LAMP1 **(F)** proteins in U266B1 treated with AUTAC (5 μM) and/or CFZ (10 nM) for 16h (scale 10 μM). Three biological replicates for each cell line were used to perform Western blotting. The uncropped blots are shown in the Supplemental Materials. Statistical significance of each condition compared to the indicated control or treatment was determined using unpaired Student’s t-test or two-way ANOVA with Tukey or Sidak post hoc tests, as appropriate. Data are represented as mean ± SEM. Significance levels are indicated as follows: * p < 0.05, ** p < 0.01, *** p < 0.001 and **** p < 0.0001.

We then extended this approach to solid tumor non-small cell lung cancer (NSCLC) cell lines NCI-H1975 and proteasome inhibitor-resistant NCI-H1975^R^. AUTAC alone demonstrated significant efficacy in degrading Mcl1, accompanied by autophagy induction, as indicated by increased LC3I/II levels (Fig. 3C). The combination of CFZ (2.5 or 10 nM) with AUTAC further enhanced Mcl1 degradation and autophagic responses compared to AUTAC alone in both cell lines (Fig. 3C).

To explore whether CFZ could amplify the Mcl1 degradation achieved with lower doses of AUTAC, we treated U266B1 cells with CFZ (10 nM) in combination with varying AUTAC doses (500 nM, 1 μM, and 5 μM). This combination resulted in greater Mcl1 degradation compared to each dose of AUTAC alone, along with increased autophagic flux confirmed by LC3I/II accumulation (Fig. 3D).

Finally, we validated CFZ’s ability to enhance Mcl1 degradation by AUTAC using immunofluorescence staining in U266B1 cells treated with AUTAC and/or CFZ for 16 hours. While AUTAC alone reduced Mcl1 levels, the combination with CFZ led to a more pronounced reduction (Fig. 3E and F). Both AUTAC and CFZ alone increased LC3B and the lysosomal marker LAMP1 levels, indicative of active autophagic flux; however, the combination treatment resulted in significant accumulation of these markers. Increased co-localization of Mcl1 with LC3B and LAMP1 was observed in U266B1 cells treated with AUTAC alone or in combination with CFZ (Fig. 3E and F). Taken together, our findings establish that autophagy inducers, particularly CFZ, enhance the ability of AUTAC to degrade Mcl1 via amplifying autophagic flux.

### CFZ-mediated Enhancement of AUTAC Activity is Dependent on NRF1, p62/SQSTM1 and Functional Autophagy

We have previously demonstrated that NRF1 is a key regulator of cellular responses to proteotoxic stress induced by proteasome inhibition ^68–71^. NRF1 facilitates the upregulation of proteasome genes to restore proteasome activity and promotes autophagy, providing cells with alternative pathways to degrade ubiquitylated protein aggregates. These mechanisms enable cells to mitigate proteotoxic stress. Notably, our group has shown that NRF1 knockout abolishes carfilzomib-induced autophagy, resulting in the accumulation of autophagosome vacuoles in NRF1-deficient cells ^13^. Initially, we showed that AUTAC-mediated activities—including Mcl1 degradation (Fig. 4A), autophagy induction, and cell death, as evidenced by cleaved caspase 3, PARP cleavage, and reduced cell viability (Fig. 4A and B)—are not dependent on NRF1. To further validate our previous findings, we examined the effects of NRF1 knockout on CFZ–mediated transcriptional responses. NRF1 knockout abolished CFZ-induced upregulation of autophagy-related genes, including p62/SQSTM1, ATG4A, CTSD, GABARAPL1, and VPS37A, as well as proteasome-related genes, including PSMA7, PSMC4, and PSMD12 (Fig. 4C). Consistently, NRF1 knockout also suppressed CFZ-induced upregulation of autophagy-related proteins, including p62/SQSTM1, CTSD, GABARAPL1, and LAMP1 (Fig. 4D). Furthermore, NRF1 knockout abolished CFZ-mediated autophagy induction, as confirmed by autophagic flux inhibition assay using CQ (Fig. 4E).

**Figure 4.**
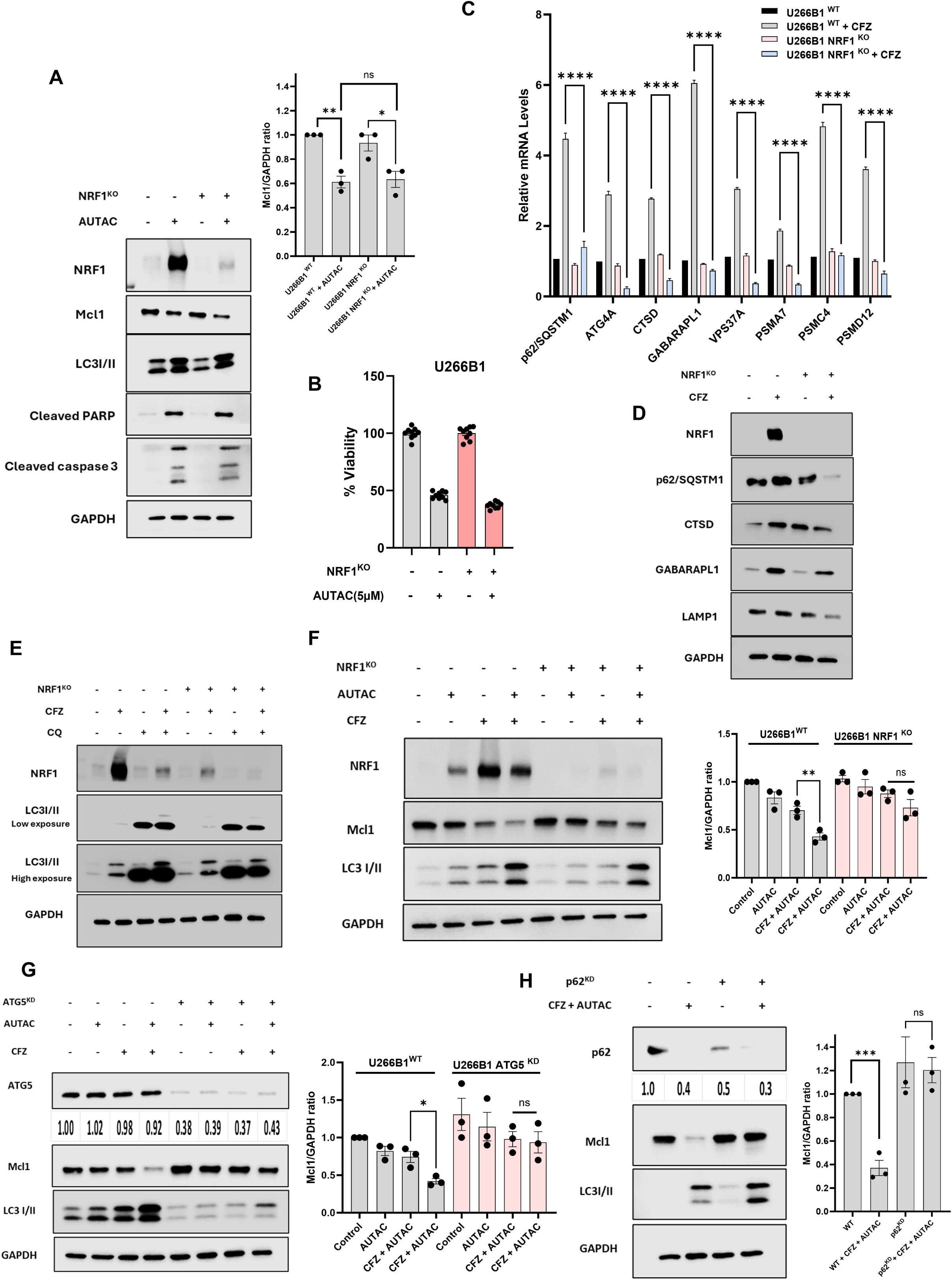
Carfilzomib-mediated potentiation of AUTAC activity is dependent on NRF1, p62/SQSTM1 and functional autophagy. **(A)** Western blot analysis for NRF1, Mcl1, cleaved PARP, cleaved caspase 3 and LC3I/II proteins in U266B1 ^WT^ and U266B1 NRF1^KO^ cell lines treated with AUTAC (5 μM) for 24h. GAPDH was used as the loading control. **(B)** The percentage of cellular viability for U266B1 ^WT^ and U266B1 NRF1^KO^ cell lines treated with AUTAC (5 μM) for 72h. **(C)** qRTPCR for the indicated genes in U266B1 ^WT^ and U266B1 NRF1^KO^ cell lines after treating with CFZ (20nM) for 20h. **(D)** Western blot analysis for NRF1, p62/SQSTM1, CTSD, GABARAPL1, and LAMP1 proteins in U266B1 ^WT^ and U266B1 NRF1^KO^ cell lines treated with CFZ (10 nM) for 16h. GAPDH was used as the loading control. **(E)** Western blot analysis for NRF1, and LC3II in U266B1 ^WT^ and U266B1 NRF1^KO^ cell lines after treating with CQ (20 μM), and/or CFZ (10 nM) for 16h. GAPDH was used as the loading control. **(F)** Western blot analysis for NRF1, Mcl1, and LC3I/II proteins in U266B1 ^WT^ and U266B1 NRF1^KO^ cell lines treated with AUTAC (5 μM) and/or CFZ (10 nM) for 16h. GAPDH was used as the loading control **(G)** Western blot analysis for ATG5, Mcl1, and LC3I/II proteins in U266B1^WT^ and U266B1 ATG5 ^KD^ cell lines treated with AUTAC (5 μM) and/or CFZ (10 nM) for 16h. GAPDH was used as the loading control. **(H)** Western blot analysis for p62/SQSTM1, Mcl1, and LC3I/II proteins in U266B1^WT^ and U266B1 p62 ^KD^ cell lines treated with AUTAC (5 μM) and/or CFZ (10 nM) for 16h. GAPDH was used as the loading control. Three biological replicates for each cell line were used to perform Western blotting. The uncropped blots are shown in the Supplemental Materials. Statistical significance of each condition compared to the indicated control or treatment was determined using unpaired Student’s t-test or two-way ANOVA with Tukey or Sidak post hoc tests, as appropriate. Data are represented as mean ± SEM. Significance levels are indicated as follows: * p < 0.05, ** p < 0.01, *** p < 0.001 and **** p < 0.0001.

We then assessed the impact of NRF1 knockout on Mcl1 degradation mediated by the combination of AUTAC and CFZ in U266B1 cells. NRF1 knockout rescued Mcl1 levels, preventing the reduction achieved by the combination of AUTAC and CFZ in wild-type cells. This effect was accompanied by diminished autophagic flux, as evidenced by decreased LC3I/II levels in NRF1-deficient cells compared to wild-type controls (Fig. 4F). To further validate the role of autophagy in this context, we employed an orthogonal approach. Knockdown of ATG5, an essential autophagy gene, significantly attenuated the Mcl1 degradation observed with the CFZ and AUTAC combination in U266B1 cells (Fig. 4G). Together, these findings underscore the pivotal role of NRF1 in CFZ-induced autophagy and highlight the necessity of functional autophagy for the synergistic activity of CFZ and AUTAC in degrading Mcl1.

We further validated the role of p62/SQSTM1 in mediating AUTAC-induced Mcl1 degradation, even in combination with CFZ. The knockdown of p62 completely abolished Mcl1 degradation induced by the combination of CFZ and AUTAC (Fig. 4H), underscoring the essential role of p62 in mediating this selective degradation pathway.

### Combination of AUTAC and Carfilzomib Overcomes Proteasome Inhibitor Resistance *In Vitro* and Suppresses Tumor Growth *In Vivo*

Chemotherapeutic agents modulate autophagy in distinct ways, including cytotoxic, cytostatic, non-protective, and cytoprotective forms ^72–74^. CFZ has been shown to induce cytoprotective autophagy ^18–20,66^, which we confirmed in our study using autophagy inhibitors like CQ and bafilomycin (BAF), which enhanced CFZ toxicity (weak synergy except in U266B1 cells with CQ) in both U266B1 cells and proteasome inhibitor-resistant U266B1^R^ cells (Supplementary Fig. 5C, D, and E). Furthermore, to substantiate the role of Mcl1 in mediating resistance to proteasome inhibitors, we generated Mcl1-overexpressing U266B1 cells (U266B1 Mcl1^OE^) and assessed their viability in response to varying concentrations of CFZ, in comparison to U266B1^WT^ and the resistant U266B1^R^ cell line (Supplementary Fig. 6A and B). U266B1 Mcl1^OE^ cells demonstrated a marked resistance to CFZ relative to their wild-type counterparts (Supplementary Fig. 6A). Additionally, western blot analysis revealed elevated Mcl1 expression levels in U266B1^R^ cells compared to parental U266B1 (Supplementary Fig. 6B), further corroborating the pivotal role of Mcl1 in conferring resistance to proteasome inhibition in multiple myeloma.

Given AUTAC’s ability to degrade Mcl1 in an autophagy-dependent manner, we hypothesized that combining AUTAC with CFZ could enhance its cytotoxic effects. Indeed, this combination reduced the viability of multiple myeloma cell lines, including U266B1, AMO-1, MM.1S, and RPMI-8226, while also sensitizing resistant lines (U266B1^R^ and AMO-1^R^) to CFZ (Fig. 5A). This reduction in cell viability occurred in a strongly synergistic manner across all tested cell lines, as confirmed by Bliss synergy scoring. Western blot analysis revealed increased cleaved caspase 3 levels, indicating apoptosis induction in both wild-type and resistant cells (Fig. 5B). We further demonstrated that AUTAC significantly potentiated CFZ-mediated cytotoxicity in U266B1 Mcl1^OE^ (Supplementary Fig. C). This effect was accompanied by a pronounced reduction in Mcl1 protein level and a marked increase in apoptotic markers, including cleaved caspase 3 and cleaved PARP (Supplementary Fig. D). To further validate the selectivity of AUTAC- and CFZ-induced cytotoxicity and to determine its dependence on p62/SQSTM1, we compared U266B1^WT^ cells with U266B1 p62^KD^ cells (Fig. 5C). Notably, AUTAC enhanced CFZ sensitivity in U266B1^WT^ cells, whereas this effect was completely abrogated in p62^KD^ cells (Fig. 5C). Collectively, these findings underscore the selective capacity of AUTAC to redirect cytoprotective autophagy toward targeted Mcl1 degradation in a p62-dependent manner, thereby amplifying apoptotic cell death.

**Figure 5.**
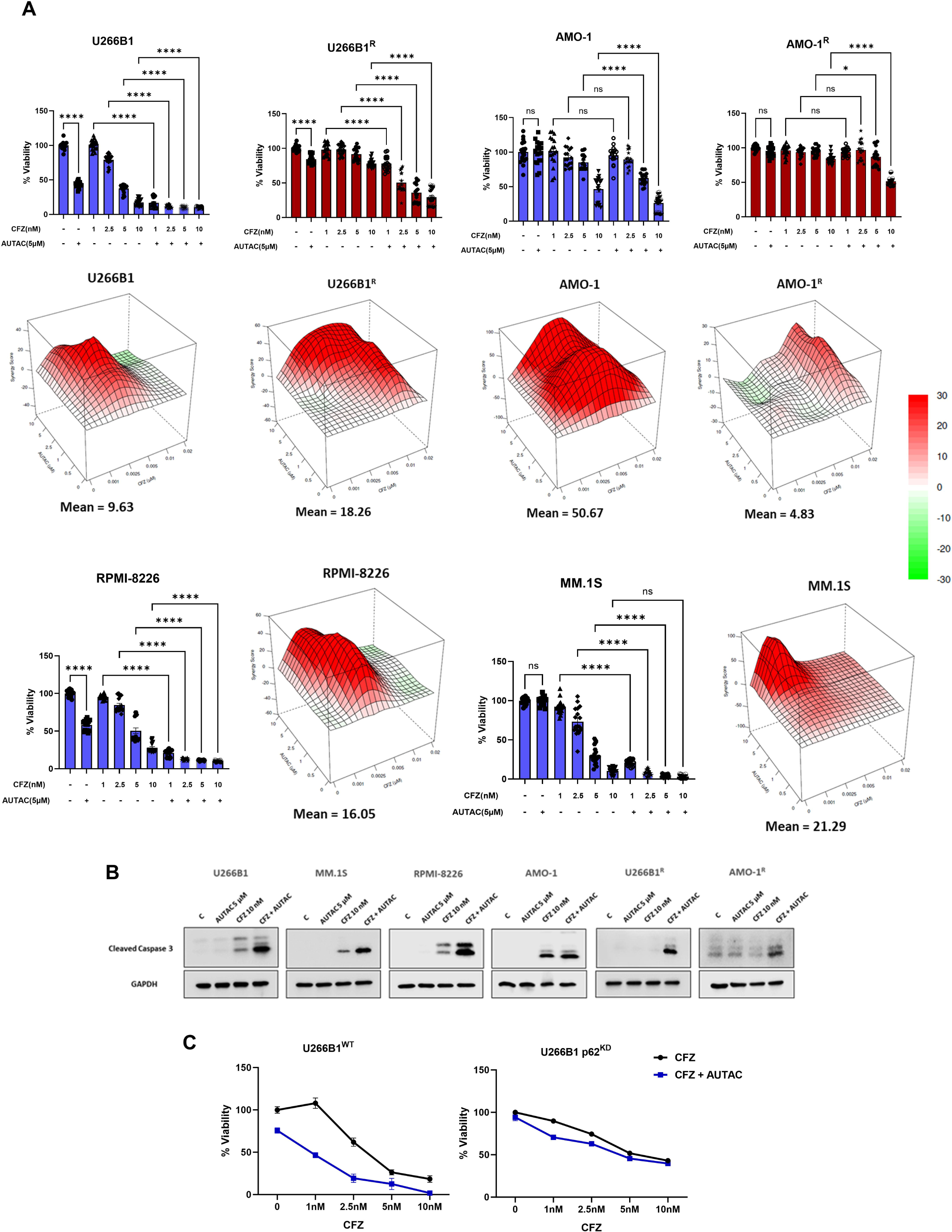
Carfilzomib in combination with AUTAC synergistically induces cell death in wild-type and proteasome inhibitor-resistant multiple myeloma cells. **(A)** The percentage of cellular viability for U266B1, MM.1S, AMO-1, RPMI-8226, U266B1^R^ and AMO-1^R^ cell lines treated with AUTAC (5 μM) for 72h, then the indicated concentrations of CFZ were added for the last 24h. The synergy between AUTAC and CFZ were determined using Bliss score. **(B)** Western blot analysis for cleaved caspase 3 protein in multiple myeloma cell lines after 16h (U266B1, MM.1S, RPMI-8226), and 24h (AMO-1) treatment with AUTAC (5 μM) and/or CFZ (10 nM), while U266B1^R^ and AMO-1^R^ cells treated for 48h with AUTAC (5 μM), then CFZ (10 nM) was added for the last 24h. GAPDH was used as the loading control. **(C)** The percentage of cellular viability for U266B1^WT^ and U266B1 p62^KD^ cell lines treated with AUTAC (5 μM) for 72h, then the indicated concentrations of CFZ were added for the last 24h. Three biological replicates for each cell line were used to perform cell viability assay and Western blotting. In each independent replicate of synergy assays, a single well was used for each drug combination. The uncropped blots are shown in the Supplemental Materials. ns indicates non-statistical significance, while * p < 0. 05, and **** p < 0.0001 indicates statistical significance of each condition compared to the indicated control/treatment as determined using two-way ANOVA with Sidak’s post hoc test.

To confirm the therapeutic superiority of the combination, we investigated its effect *in vivo* using NSG mice subcutaneously injected with U266B1 cells. Treatment with CFZ alone resulted in a significant reduction in tumor volume (Fig. 6A); however, subsequent treatment with AUTAC led to a further reduction in tumor volume compared with either treatment alone (Fig. 6A), without significant toxicity (Fig. 6B). Mechanistically, CFZ enhanced the ability of AUTAC to reduce Mcl1 protein levels, while the combination had no impact on Bclxl or Bcl2 levels (Fig. 6C). Furthermore, autophagy induction was evidenced by reduced p62/SQSTM1 levels and increased LC3II accumulation (Fig. 6B). Subsequently, we conducted a comparative analysis of AUTAC against several well-characterized Mcl1 inhibitors, including S63845, AMG 176, UMI-77, AZD-5991, and A-1210477. Given the well-documented cardiotoxicity observed with Mcl1 inhibition in clinical settings, we specifically assessed the cytotoxic profiles of these compounds in AC16 human cardiomyocytes. Although AUTAC (IC□□= 5 µM) was less potent against U266B1 cells compared to AMG176 (IC□□= 750 nM), S63845 (IC□□= 200 nM), and AZD-5991 (IC□□= 100 nM), it demonstrated a more favorable safety profile. Notably, AUTAC exhibited no detectable cardiotoxicity in this model. In contrast, S63845 (AC16 IC□□= 100 nM), AMG176 (AC16 IC □□= 2.5 µM), and AZD-5991 (AC16 IC□□ = 100 nM) induced pronounced cytotoxic effects (Fig. 5D). These findings highlight the potentially superior cardiac safety profile of AUTAC. Next, we evaluated the synergistic potential of these Mcl1 inhibitors in combination with CFZ in both U266B1 and its resistant counterpart, U266B1R (Fig. 7). Although AUTAC, along with UMI-77 and A-1210477, demonstrated comparatively lower standalone potency relative to S63845, AMG 176, and AZD-5991, AUTAC exhibited the highest degree of synergy with CFZ across both cell lines, as quantified by Bliss synergy scores (Fig. 7). In contrast, the other inhibitors displayed variable interactions ranging from weak synergy, as observed with A-1210477, to merely additive effects or even antagonism with to S63845, UMI-77, AMG 176, and AZD-5991 (Fig. 7), in consistent with several reports ^75–78^. These findings underscore the unique advantage of AUTAC in harnessing CFZ-induced autophagic flux to enhance Mcl1 degradation, thereby potentiating its combinatorial efficacy.

**Figure 6.**
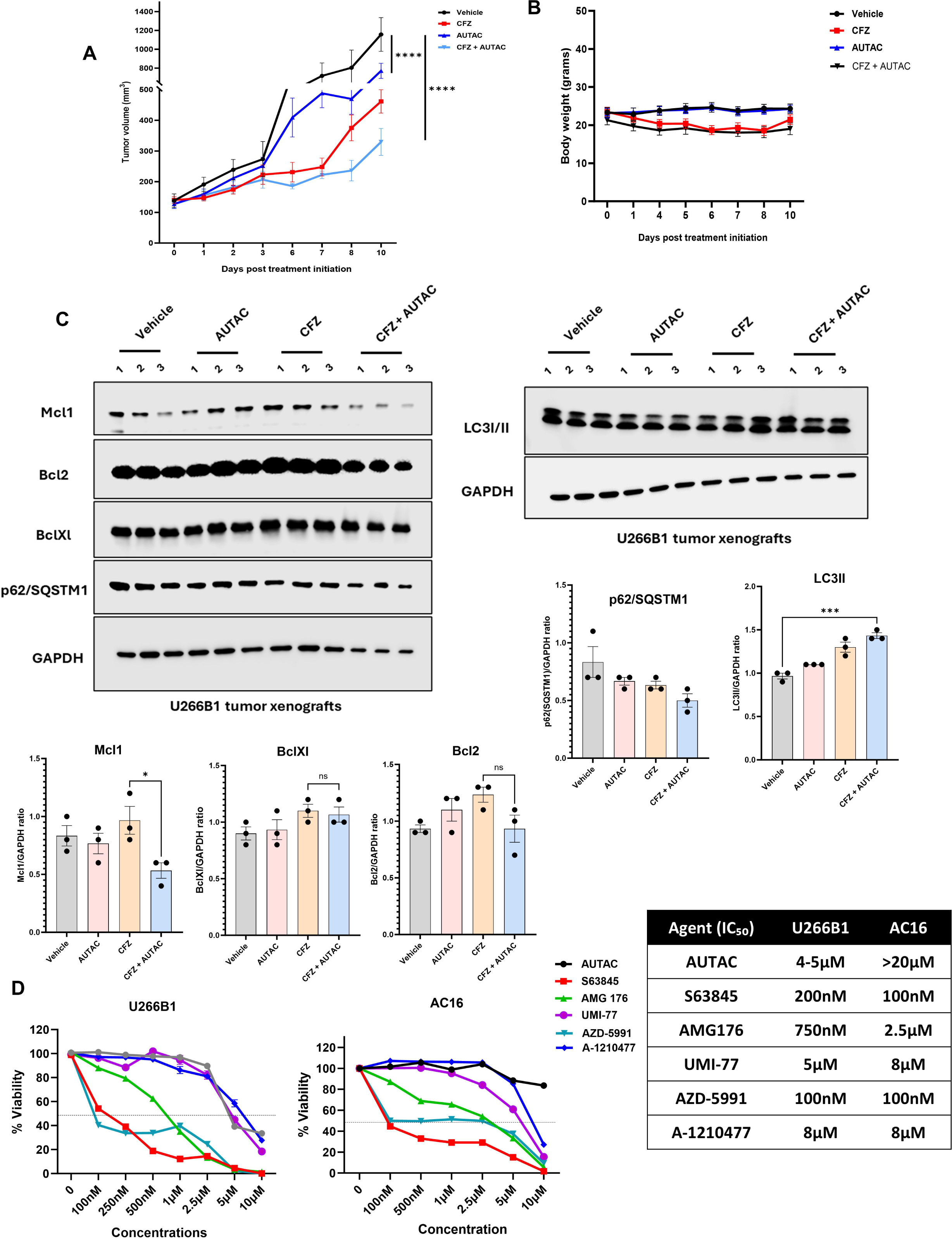
AUTAC potentiates the *in vivo* antitumor activity of CFZ in U266B1 xenograft models. **(A)** Tumor volumes of U266B1 xenografts following treatment with vehicle, CFZ (5 mg/kg), and/or AUTAC (50 mg/kg). CFZ was administered on days 1, 4, and 7, while AUTAC treatment was initiated on day 7 and continued daily. **(B)** Body weights of mice bearing U266B1 xenografts during treatment with vehicle, CFZ (5 mg/kg), and/or AUTAC (50 mg/kg). **(C)** Western blot analysis for Mcl1, Bcl2, Bclxl, p62/SQSTM1, and LC3II for U266B1 tumor xenografts from mice treated with vehicle, CFZ (5 mg/kg), and/or AUTAC (50 mg/kg). **(D)** The cell viability of U266B1 and AC16 cardiomyocytes treated with the indicated concentrations of AUTAC, S63845, AMG 176, UMI-77, AZD-5991, and A-1210477 for 72h with corresponding IC values calculated for each cell line. Three biological replicates for each cell line were used to perform Western blotting. The uncropped blots are shown in the Supplemental Materials. Statistical significance of each condition compared to the indicated control or treatment was determined using unpaired Student’s t-test or two-way ANOVA with Tukey or Sidak post hoc tests, as appropriate. Data are represented as mean ± SEM. Significance levels are indicated as follows: * p < 0.05, ** p < 0.01, *** p < 0.001 and **** p < 0.0001.

**Figure 7.**
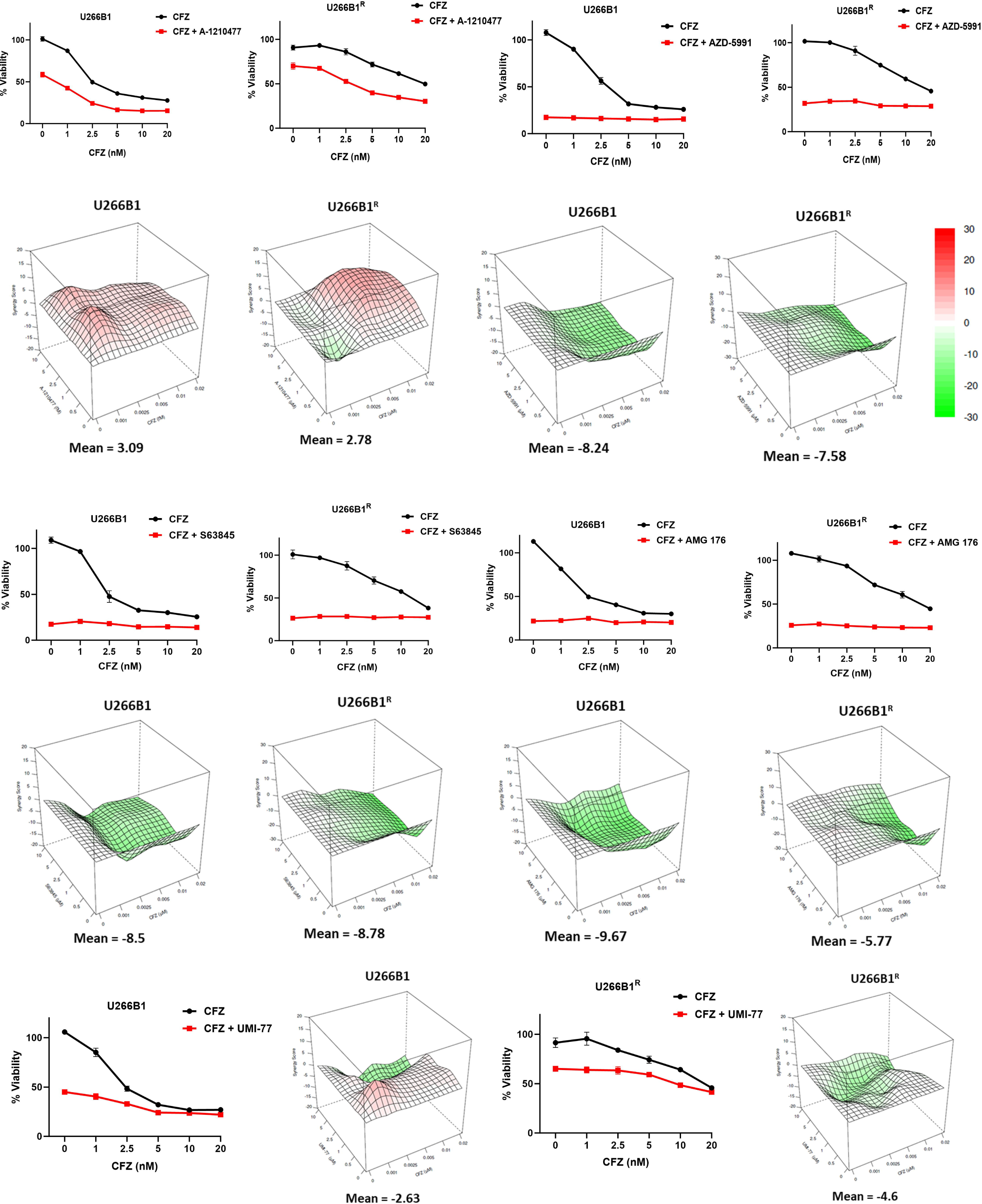
Classical Mcl1 inhibitors do not synergize with carfilzomib in inducing cell death in wild-type or proteasome inhibitor-resistant multiple myeloma cells. **(A)** The percentage of cellular viability for U266B1, and U266B1R cell lines treated with 5μM of S63845, AMG 176, UMI-77, AZD-5991, and A-1210477 for 72h, then the indicated concentrations of CFZ were added for the last 24h. The synergy between AUTAC and CFZ were determined using Bliss score. Three biological replicates for each cell line were used to perform cell viability assay. In each independent replicate of the synergy assay, a single well was used for each drug combination.

In solid tumors, including non-small cell lung cancer (NSCLC), proteasome inhibitors (PIs) have shown limited efficacy ^6–8^. To improve PI efficacy in this context, we evaluated the combination of AUTAC and CFZ in wild-type (NCI-H1975) and bortezomib-resistant (NCI-H1975^R^) NSCLC cells. The combination significantly reduced cell viability compared to either treatment alone (Fig. 8A) and increased cleaved caspase 3 accumulation, confirming apoptosis (Fig. 8B). This reduction in cell viability occurred in a strongly synergistic manner in both cell lines, as confirmed by Bliss synergy scoring (Fig. 8A). Colony formation assays demonstrated that while AUTAC or CFZ alone moderately reduced the colony-forming ability of NCI-H1975 cells, their combination had a significantly stronger effect. In NCI-H1975^R^ cells, which showed minimal response to CFZ alone, the combination markedly inhibited colony formation (Fig. 8C). These findings highlight the potential of AUTAC to enhance PI efficacy by disrupting cytoprotective autophagy and promoting apoptotic cell death in both hematologic and solid tumors.

**Figure 8.**
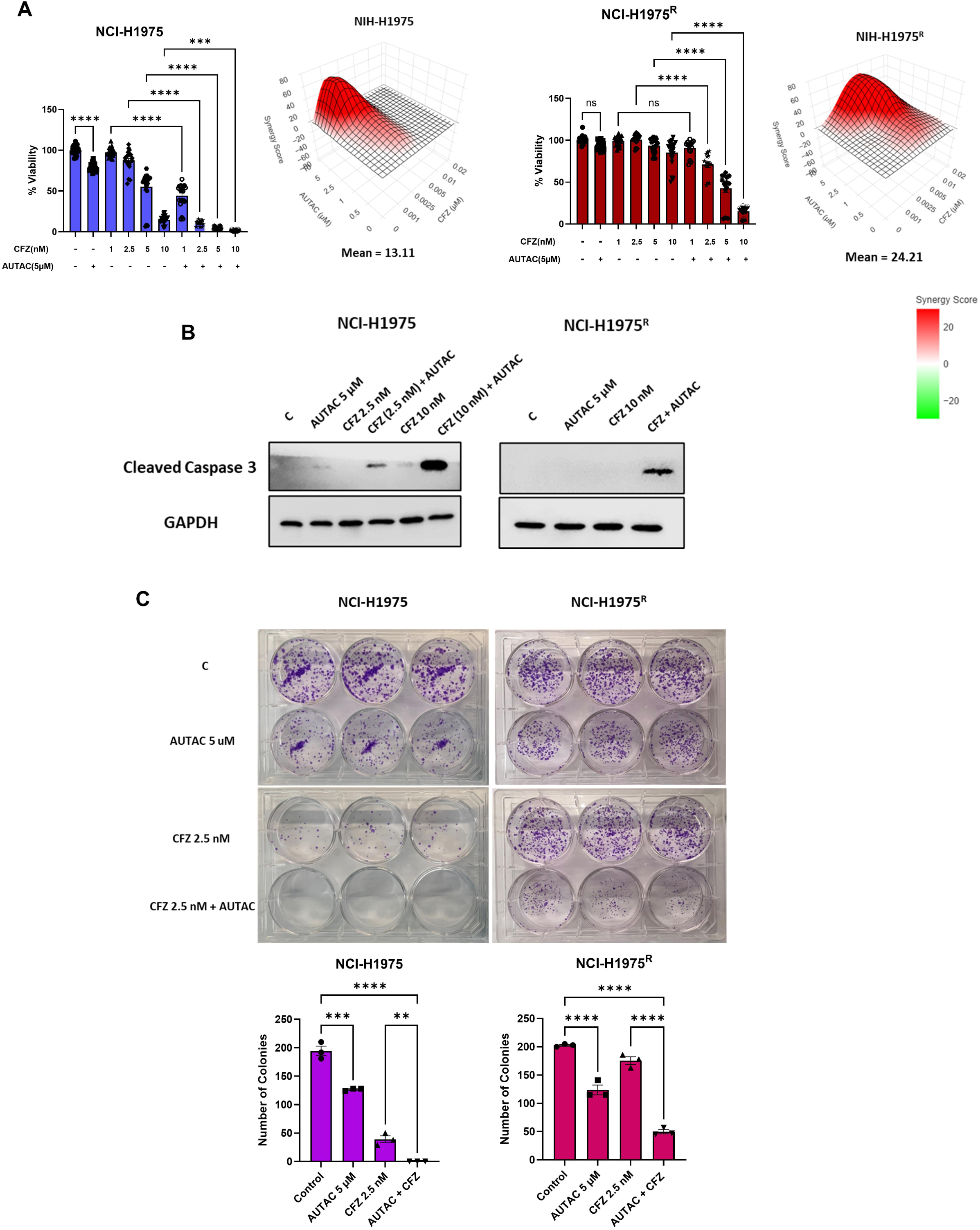
The impact of combining carfilzomib with AUTAC in wild-type and proteasome inhibitor-resistant non-small cell lung cancer cells. **(A)** The percentage of cellular viability for NCI-H1975 and NCI-H1975^R^ cells treated with AUTAC (5 μM) and/or the indicated concentrations of CFZ for 72h. The synergy between AUTAC and CFZ were determined using Bliss score. **(B)** Western blot analysis for cleaved caspase 3 protein in NCI-H1975 and NCI-H1975^R^ cells treated with AUTAC (5 μM) and/or CFZ (2.5, and 10 nM) for 72h. GAPDH was used as the loading control. **(C)** Colony formation assay and its quantification for NCI-H1975 and NCI-H1975^R^ cells treated with AUTAC (5 μM) and/or CFZ (2.5, and 10 nM) for 72h, then supplemented with media every three days for total 9 days. Three biological replicates for each cell line were used to perform cell viability assay, colony formation assay and Western blotting. In each independent replicate of the synergy assay, a single well was used for each drug combination. The uncropped blots are shown in the Supplemental Materials. ns indicates non-statistical significance, while ** p < 0.01, *** p < 0.001 and **** p < 0.0001 indicates statistical significance of each condition compared to the indicated control/treatment as determined using two-way ANOVA with Sidak’s post hoc test.

## DISCUSSION

Proteasome inhibitors (PIs) have transformed the treatment landscape for multiple myeloma, eliciting robust initial responses and significantly improving patient outcomes and survival rates. However, disease relapse and acquired resistance remain major clinical challenges ^79^. Several adaptive mechanisms could underlie resistance to PIs, including NRF1-driven autophagy and the antiapoptotic role of Mcl1 ^13–15,26–28^. In this study, we demonstrated that leveraging Mcl1-targeting AUTAC enhances the efficacy of CFZ in both hematologic malignancies, both *in vitro* and *in vivo*, and solid tumors, including proteasome inhibitor-resistant models.

The concept of AUTAC originates from studies on selective autophagy triggered by infection, where nitric oxide (NO) and ROS mediate the degradation of bacterial proteins. Specifically, NO facilitates the conversion of cytoplasmic cyclic guanosine monophosphate (cGMP) to 8-nitro-cGMP, which modifies bacterial surface proteins via S-guanylation, targeting them for autophagic degradation ^22,34,80–82^. Takahashi et al. ^23^ harnessed this pathway to develop the first and second generations of AUTACs, substituting cysteine with a pyrazole ring in the degradation tag. Building on this framework, we developed a second-generation AUTAC to selectively degrade Mcl1. Our results demonstrate that AUTAC selectively targets Mcl1 for degradation among members of the Bcl2 protein family. Given that AUTAC induces the autophagic machinery, we confirmed that this effect is not attributable to nonspecific autophagic flux. Moreover, our data demonstrates that AUTAC activates autophagy in a Beclin1-dependent manner, rather than through mTOR suppression, nonspecific global autophagy induction, or indiscriminate ROS generation. These findings are consistent with previous reports showing that several Mcl1 inhibitors disrupt the interaction between Mcl1 and Beclin1, thereby promoting Beclin1-mediated autophagy ^54,55,83,84^. Critically, we show that the intact AUTAC molecule is required for selective Mcl1 degradation and autophagy induction, as neither the Mcl1-binding ligand nor the FBng tag alone was sufficient to elicit these effects. We further confirmed the autophagy-dependent mechanism of Mcl1 degradation by AUTAC, validated pharmacologically using early and late autophagy inhibitors (3-MA and CQ, respectively) and genetically via ATG5 knockdown.

The AUTAC technology offers distinct advantages over ubiquitin-proteasome system (UPS)-based degradation strategies ^23,85^. Unlike UPS-targeting approaches, which are limited to soluble proteins, AUTAC has the potential to target a broader range of cellular cargoes, including non-proteinaceous biomolecules and organelles such as mitochondria ^23,85^. In terms of kinetics, we observed initial signs of AUTAC-mediated protein degradation as early as 3 hours post-treatment (data not shown), with degradation efficiency increasing over time. This gradual onset is likely reflective of the more complex, multistep nature of autophagy compared to the relatively direct mechanism of UPS-mediated degradation ^23,85^. Mechanistically, AUTAC exerts its degradative function through K63-linked polyubiquitination of target proteins, as first demonstrated by Takahashi et al.^23^. In their study, the degradation activity of AUTAC was abrogated upon genetic elimination of p62/SQSTM1, highlighting the essential role of this selective autophagy receptor in the pathway. In line with these findings, our data confirm that AUTAC promotes the colocalization of Mcl1 with K63-linked ubiquitin chains, p62/SQSTM1, and the autophagosomal marker LC3B, suggesting that Mcl1 is recruited into autophagosomes via the canonical selective autophagy machinery. We further corroborated the colocalization findings by Co-IP analysis, which demonstrated that AUTAC treatment enhanced the association of K63-linked ubiquitin and p62/SQSTM1 with Mcl1. Furthermore, shRNA-mediated depletion of p62 abolished the AUTAC-induced degradation of Mcl1, further substantiating the requirement of this receptor in mediating cargo selectivity. The role of p62/SQSTM1 is particularly significant, as it serves as a molecular bridge linking ubiquitinated substrates to the autophagosome ^57–60^. It binds K63-polyubiquitinated proteins via its C-terminal ubiquitin-associated (UBA) domain and facilitates their sequestration into autophagosomes through direct interaction with LC3 via its LC3-interacting region (LIR) ^57–60^. This mechanism reinforces the specificity of AUTAC in targeting distinct proteins, such as Mcl1, for autophagic degradation.

Interestingly, 8-nitro-cGMP has been reported to act as a molecular tag that facilitates K63-linked polyubiquitination ^22^. In this context, Yamada et al.^86^ demonstrated that the FBXO2-SCF E3 ubiquitin ligase complex recognizes bacterial surface glycans modified with 8-nitro-cGMP, thereby inducing K63-linked polyubiquitination and initiating xenophagic clearance. Therefore, we investigated the specific E3 and E2 ligases responsible for mediating AUTAC-induced K63-linked ubiquitination of Mcl1. Senichkin et al.^87^ reported that the E3 ubiquitin ligase TRAF6 mediates K63-linked ubiquitination of Mcl1, resulting in its stabilization rather than proteasomal degradation, which also reported by Sancho et al. ^88^. In contrast, proteasomal turnover of Mcl1 is primarily driven by K48-linked ubiquitination, largely mediated by the E3 ligase Mule (also known as HUWE1)^87,89^. Similarly, Choi et al. ^61^ demonstrated that the viral protein HTLV-1 Tax promotes non-degradative K63-linked ubiquitination of Mcl1, thereby enhancing its stability. These reports confirm that Mcl1 is subjected to K63 ubiquitination, which mediate Mcl1 stabilization and involved in several signaling purposes^88,90–92^. However, under stress conditions, K63-linked ubiquitination can function as an autophagic signal, by promoting recognition by selective autophagy receptors ^92–94^. In parallel, our findings demonstrate that AUTAC engages TRAF6 together with UBC13, the principal E2 ubiquitin-conjugating enzyme responsible for K63 chain assembly ^95–97^, to promote K63-linked ubiquitination of Mcl1. These results reveal a model in which AUTAC selectively harnesses the TRAF6–UBC13 axis to redirect Mcl1 toward degradation, through K63 ubiquitination and subsequent recognition by p62/SQSTM1, thereby reinforcing the mechanistic specificity underlying AUTAC-mediated targeting. Additionally, AUTAC markedly reduced the viability of multiple myeloma and NSCLC cells through robust induction of apoptosis. In contrast, genetic silencing of p62, TRAF6, or UBC13 effectively abolished AUTAC-mediated cytotoxicity, demonstrating that its anti-tumor activity is strictly dependent on the autophagic ubiquitin-tagging pathway and excluding the possibility of off-target effects.

We further exploited the cytoprotective autophagy induced by CFZ, repurposing it as a degradation mechanism for Mcl1. The combination of CFZ and AUTAC markedly reduced Mcl1 levels in both wild-type and proteasome inhibitor-resistant multiple myeloma and NSCLC cells, accompanied by a robust upregulation of autophagic flux. Importantly, NRF1, p62/SQSTM1 and functional autophagy were essential for Mcl1 degradation, as shown through genetic inhibition approaches. These mechanistic insights translated into significant reductions in cellular viability and increased apoptosis, highlighting the ability of AUTAC to sensitize resistant cells to CFZ. We further demonstrated that the combination of AUTAC and CFZ significantly suppressed U266B1 tumor xenografts growth compared with either monotherapy alone. These findings provide a proof of concept that combining proteasome inhibitors can potentiate the degradative capacity of AUTAC technology, suggesting its broader applicability to targets beyond Mcl1. The interplay between autophagy and PI resistance is well-established ^18–20,66^, and we confirmed that autophagy inhibition using CQ or BAF augmented CFZ-induced cytotoxicity. However, the lack of selective autophagy inhibitors and the adverse effects of existing agents, such as CQ and HCQ, limit their clinical utility ^98–101^. This study introduces a novel strategy of harnessing PIs’ cytoprotective autophagy, rather than inhibiting it, by targeting resistance-associated proteins like Mcl1 for autophagic degradation.

We also evaluated the efficacy and safety profile of AUTAC in comparison with several classical Mcl1 inhibitors. Among all agents tested, AUTAC exhibited the lowest cytotoxicity toward cardiomyocytes while demonstrating the most pronounced synergistic effect with CFZ. Given the well-documented cardiotoxicity associated with proteasome inhibitor therapy, our findings suggest that AUTAC may serve as a valuable adjunct to enhance the antitumor efficacy of proteasome inhibitors while minimizing their dose-limiting toxicities. This combinatorial approach holds promise for reducing the required dosage of proteasome inhibitors, thereby preserving therapeutic benefit while minimizing off-target adverse effects, particularly in cardiac tissue. Further research is necessary for the structural optimization of AUTAC to enhance its potency and therapeutic efficacy.

## MATERIALS AND METHODS

### Cell Culture

MM.1S (CRL-2974; ATCC), RPMI-8226 (CCL-155; ATCC), AMO-1, AMO-1 resistant/ AMO-1^R^ (both sensitive and resistant cell lines were obtained from Dr. Bianchi Lab, Division of Hematology, Brigham and Women’s Hospital, Harvard Medical School), NCI-H1975 ^102^, NCI-H1975 resistant/ NCI-H1975^R^ (NCI-H1975 were continuously exposed to increasing concentrations of bortezomib (up to 10 nM) over a period of 10 weeks, with surviving cells maintained in 10 nM), U266B1 resistant/ U266B1^R^ ^103^, U266B1 (TIB-196; ATCC), U266B1 ATG5^KD^ ^104^, U266B1 p62^KD104^, and their derivatives were grown in Roswell Park Memorial Institute Medium (RPMI) (Gibco), while AC16 cells (CRL-3568; ATCC) were grown in Dulbecco’s Modified Eagle Medium (DMEM; Gibco). Both media were supplemented with 10% fetal bovine serum (FBS; Atlanta Biologicals), 1X penicillin-streptomycin (Pen/Strep; Invitrogen) and 5 µg/ml Plasmocin Prophylactic (PP; InvivoGen). All cell lines were grown at 37°C in a humidified incubator with 5% CO2.

### Cell line provenance and quality control

Cell lines obtained from ATCC were used according to supplier documentation. Cell lines obtained from external laboratories were used as received. Cell lines were authenticated by the supplier. Mycoplasma contamination was monitored routinely using InvivoGen’s MycoStrip® detection assay and prevented by Plasmocin prophylaxis.

### Study Design, Replicates, and Sample Size

Unless otherwise indicated, all *in vitro* experiments were performed with at least three independent biological replicates. Within each independent experiment, assays were performed in technical triplicate, except for cell viability assays, which were performed with six technical replicates per condition.

### Sample size justification

Sample sizes for *in vitro* experiments were selected based on prior experience with these assays and expected effect sizes from preliminary work. No formal a priori power calculations were performed for *in vitro* studies.

For animal studies, n values are stated explicitly (see Mouse xenograft model). Group sizes were selected based on prior xenograft experience and expected effect sizes; no formal power calculation was performed.

### Inclusion/Exclusion Criteria and Data Exclusion

No animals or biological replicates were excluded from analysis. For *in vitro* studies, experiments were repeated only in the case of clear technical failure, and these repeats were not based on outcome. Criteria were applied consistently and were not modified post hoc.

### Randomization and Blinding. Randomization (animal studies)

Xenograft implantation, dosing, and tumor measurements were performed by the Massey Cancer Center Mouse Core. Once tumors reached 75–100 mm³, mice were randomized into treatment groups using a randomization procedure implemented by the core facility. Baseline tumor volumes did not differ significantly between groups prior to treatment initiation (TV, P = 0.990), confirming balanced allocation. Randomization was performed prior to initiation of drug treatment.

### Randomization (*in vitro*)

For plate-based assays, wells were assigned to conditions without systematic bias.

### Blinding

Investigators were not routinely blinded to treatment allocation during *in vitro* experiments. Xenograft dosing and tumor measurements were conducted by the Massey Cancer Center Mouse Core. Blinding during tumor measurement was performed according to standard core operating procedures when feasible. The study investigators did not participate in tumor measurements. Quantitative analyses (western blot densitometry, fluorescence intensity measurements, and colocalization analyses) were conducted using predefined ImageJ analysis parameters applied uniformly across all conditions to minimize subjective bias.

### Western blotting

Cells were plated and treated in six-well plates, then collected at the indicated time points and washed once with ice-cold PBS. Cells lysed using RIPA Buffer (20 mM Tris-HCl pH 7.4, 1% sodium deoxycholate, 1% Triton X-100, 0.1% SDS, 150 mM NaCl, and 1 mM EDTA) containing protease and phosphatase Inhibitor cocktail (Cat No: PI78447; Thermo Fisher Scientific), then incubated on ice for 15 min, and centrifuged at 14,000 rpm for 10 min at 4°C. Protein concentrations were measured by the bicinchoninic acid (BCA) assay. Western blot samples were prepared with 4X Laemmli protein sample buffer (Cat No:1610747; Bio-Rad) with the addition of 2-mercaptoethanol on the day of lysis, followed by boiling for 7 min. 10-15 µg of protein was used for SDS-PAGE, gels were transferred onto Immobilon FL PVDF membranes (Cat No: IPVH00010; Millipore Sigma). Membranes were blocked with 5% blotting grade blocker (Cat No: 1706404; BIO-RAD) in TBST for 1-2 h.

Blots were incubated with the indicated primary antibodies overnight, then washed with TBST and incubated with secondary antibodies for 1 h at room temperature. The antibodies used were LC3B (Cat No:2775S; Cell Signaling) at 1:1,000, NRF1 (Cat No:8052S; Cell Signaling) at 1:1,000, cleaved caspase-3 (Cat No:9661L; Cell Signaling) at 1:500, cleaved PARP (Cat No: 5625S; Cell Signaling) at 1:1000, ATG5 (Cat No: 12994S; Cell Signaling) at 1:1000, SQSTM1/p62 (Cat No:5114S; Cell Signaling) at 1:1000, Mcl1 (Cat No: 5453S; Cell Signaling) at 1:500, Bcl2 (Cat No: 3498S; Cell Signaling) at 1:1,000, Bclxl (Cat No: 2764 S; Cell Signaling) at 1:1,000, TRAF6 (Cat No: 8028S; Cell Signaling) at 1:1,000, UBC13 (Cat No: 4919S; Cell Signaling) at 1:1,000, Beclin1 (Cat No: 3495S; Cell Signaling) at 1:1,000, p-Beclin1 (Ser15) 1:500 (Cat No: D4B7R; Cell Signaling), K63 1:1000 (Cat No: HWA4C4; Invitrogen) and GAPDH (Cat No: 5174S; Cell Signaling) at 1:10,000. Goat anti-Rabbit Igg HRP conjugate antibody (Cat No:1706515; BIO-RAD) at 1:10,000 dilutions. Blots then washed with TBST, then incubated with clarity western ECL substrate (Cat No: 1705060; BIO-RAD, or Cat No: 1705062; BIO-RAD) for 5 minutes. Imaging of the blots was done by the LiCOR Odyssey Fc Imaging System. In all pharmacologic and genetic manipulation experiments, appropriate matched controls were included. DMSO served as the vehicle control for compound treatments. GFP-expressing control cells (shGFP or KO-GFP) were used as negative controls for knockdown and knockout studies, respectively.

Western blot densitometric analysis was performed using ImageJ software. Band intensities were quantified from at least three independent experiments. Protein expression levels were normalized to the corresponding GAPDH loading control and subsequently expressed relative to the control group.

### Co-immunoprecipitation (Co-IP)

U266B1 cells were seeded in 6-well plates and treated with AUTAC (10 μM) for 16 h (for K63-linked ubiquitin and p62) or 24 h (for Beclin1). Cells were harvested by centrifugation and lysed in immunoprecipitation (IP) buffer (50 mM Tris-HCl, pH 7.4, 100 mM NaCl, 1 mM EDTA, 1% Triton X-100) supplemented with protease and phosphatase inhibitor cocktails (Thermo Fisher Scientific, Pierce). Lysates were incubated on ice for 10–15 min and clarified by centrifugation at 14,000 rpm for 10 min at 4°C. Protein concentration was determined using a BCA assay. All samples were normalized to equal protein concentrations and adjusted to a final volume of 300 µL with IP buffer. An aliquot of 25 µL was retained as input (total cell lysate). The remaining supernatants were subjected to immunoprecipitation using Protein A/G agarose beads and incubated with anti-Mcl1 antibody (1:50; D2W9E, Cell Signaling Technology). Beads were pre-blocked with 5% BSA for 1 h prior to incubation with lysates. After immunoprecipitation, beads were washed three times with IP buffer and proteins were eluted in Laemmli sample buffer. Eluted proteins and input samples were resolved by SDS–PAGE and analyzed by western blotting as described previously. DMSO served as the vehicle control for compound treatments.

### Quantitative real-time PCR (qRT–PCR)

U266B1^WT^ and U266B1 NRF1^KO^ cells were seeded in 6-well plates and treated with CFZ (20 nM) for 20 h. Following treatment, cells were washed once with phosphate-buffered saline (PBS) and harvested by gentle scraping in 1 mL PBS. Total RNA was isolated using the RNeasy Mini Kit (Qiagen) with on-column DNase I digestion to remove genomic DNA contamination. Complementary DNA (cDNA) was synthesized from 1 µg of total RNA using iScript Reverse Transcription Supermix (Bio-Rad) according to the manufacturer’s instructions. Quantitative real-time PCR was performed using iTaq Universal SYBR Green Supermix (Bio-Rad) on a C1000 Touch Thermal Cycler (Bio-Rad). Transcript levels were normalized to 18S rRNA as the internal reference gene. Relative gene expression was calculated using the ΔΔCt method and analyzed using CFX Manager software (v3.1; Bio-Rad). In all pharmacologic and genetic manipulation experiments, appropriate matched controls were included. DMSO served as the vehicle control for compound treatments. GFP-expressing control cells (KO-GFP) were used as negative controls for knockout studies.

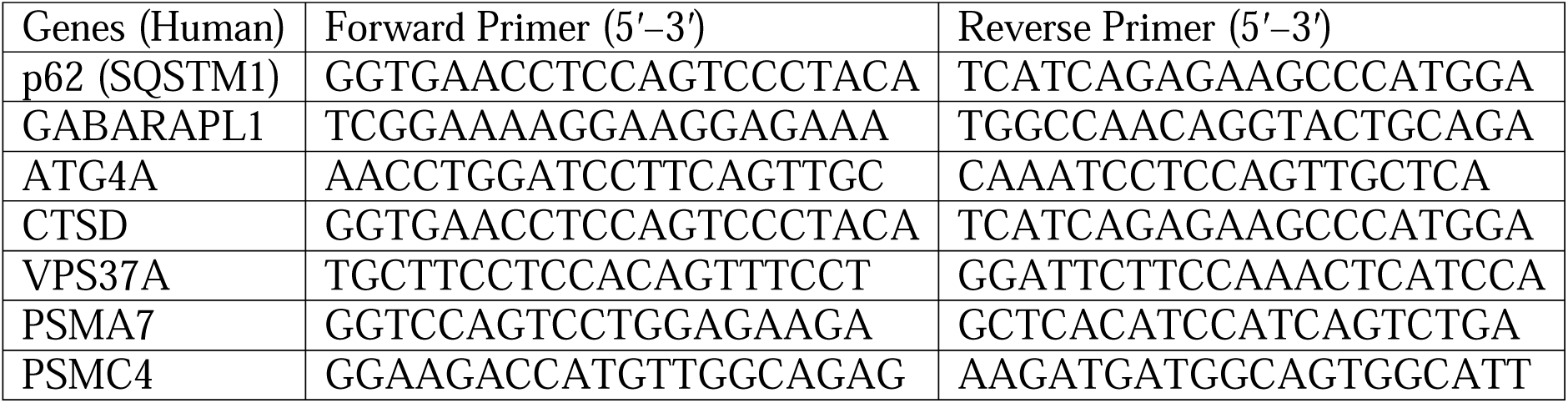
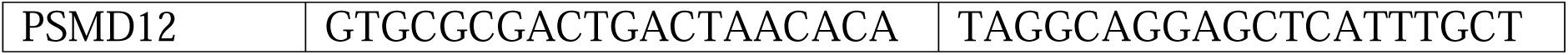

### Mouse xenograft model

All animal studies were approved by the Virginia Commonwealth University (VCU) Institutional Animal Care and Use Committee (IACUC) (Protocol AD10001417) and conducted in accordance with institutional guidelines and regulations. Animal experiments are reported in compliance with the ARRIVE 2.0 guidelines. Male and female NOD-SCID-IL2Rγ (NSG) mice (4–6 weeks old) were inoculated subcutaneously (unilaterally, right flank) with U266B1 (8 × 10^6^ cells; n =4 mice per treatment group) resuspended in Matrigel in a total volume of 100 µL. Tumor growth was monitored biweekly by manual caliper measurements, and tumor volumes were calculated using the modified ellipsoid formula ^105^: V = (L × W²)/2, where L is the longest diameter and W is the perpendicular diameter. Once tumors reached approximately 75–100 mm³, mice were randomized into treatment groups and tumor volumes were measured daily. Carfilzomib (5 mg/kg) or an equal volume of vehicle (Captisol) was administered via tail vein injection on days 1, 4, and 7. This was followed by once-daily intraperitoneal injections of vehicle (10% DMSO, 40% PEG300, 5% Tween-80, and 45% saline) or AUTAC (50 mg/kg). Tumors were harvested at the study endpoint (tumor volume of ∼1500 mm³) for downstream analyses. No animals were excluded.

### Cell viability

The CellTiter-Glo assay was utilized as the screening protocol for drugs action. Cells were plated in 96 well plate at a density of 2500-6000 cells per well and then were treated with the indicated conditions. The CellTiter-Glo assay (G7570; Promega) was performed according to standard protocol and luminescence was quantified using a POLARStar Optima cell microplate reader. Unless otherwise indicated, technical replicates: 6 wells per condition per experiment. Biological replicates: ≥3 independent experiments. The figures were generated using GraphPad Prism software. Bliss score and synergy figures were generated using synergy finder ^+^^106^. DMSO served as the vehicle control for compound treatments. GFP-expressing control cells (shGFP) were used as negative controls for knockdown.

### Cell death

The percentage of dead cells were calculated using trypan blue exclusion assay ^107^. U266B1 cells were treated with AUTAC 5 μM for 48 and 72h. Cells were stained with trypan blue, then counted. Experiments were repeated independently ≥3 times. The figures were generated using GraphPad Prism software. DMSO served as the vehicle control for compound treatments.

### Clonogenic assay

Cells plated in 6 well-plates at density of 2500-3000 cells per well, then allowed to adhere overnight. Cells treated with AUTAC 5 μM and/or CFZ (2.5-10 nM) for 72h, then drug removed, and cells replenished with fresh media every 3 days for total of 9 days. Plates were fixed with methanol, stained with crystal violet, and manually counted. Experiments were repeated independently ≥3 times. The figures were generated using GraphPad Prism software. DMSO served as the vehicle control for compound treatments.

### Lysotracker imaging

Cells were plated in 6 well plates and starved (treated with FBS depleted media) or treated with CFZ 10nM for 24h. Cells stained with DAPI (Cat No: PI62249; ThermoFisher) and Lysotracker (Cat No: L7528; ThermoFisher) stains according to the manufacture protocol. Fluorescence images of cells were taken using a brightfield inverted microscope (20X objective, Q-Color3™ Olympus Camera; Olympus, Tokyo, Japan). The merged images and quantification were generated using Image J software. The figures were generated using GraphPad Prism software. DMSO served as the vehicle control for compound treatments.

### U266B1 NRF1^KO^ cells

Edit-R human NRF1 all-in-one lentivirus sgRNA (Catalog#: SGSH12178-249373759) and Edit-R all-in-one lentiviral sgRNA hEF1a-EGFP non-targeting control (Catalog#: GSGC12062) were purchased from Horizon. For generating lentiviral supernatant, HEX 293 LXT cells were grown to 80% confluency in a 10 cm dish and transfected with lipofectamine (L3000015, ThermoFisher Scientific) with 8 µg lentivirus construct, 6 µg psPAX2 (packaging plasmid, 12260, Addgene), 4 µg pmD2.G (envelope plasmid, 12259, Addgene). Cells were incubated in OptiMEM (31985070, Thermo Fisher Scientific) transfection reagent overnight at 37 °C. The OptiMEM was replaced with fresh 5 mL 5% FBS full media, and cells were incubated at 10% CO_2_. 48 hours post-transfection, the supernatant containing virus was harvested and immediately filtered to remove cell debris. U266B1 cells were transduced with lentiviral supernatant supplemented with fresh growth media and 8 μg/mL polybrene. Successfully transduced cells were then selected by fluorescence-activated cell sorting (FACS). To confirm NRF1 knockout in U266B1 cells, WB was performed. U266B1 and U266B1 NRF1^KO^ cells were treated with 10 nM CFZ for 6 hours.

### TRAF6,UBC13, and Mcl1 Knockdown

Knockdown was achieved using smart pool (Dharmacon) of small interfering RNAs (siRNAs). For TRAF6, GGAGAAACCUGUUGUGAUU, GGACAAAGUUGCUGAAAUC, GUUCAUAGUUUGAGCGUUA, and GAUAUGAUGUAGAGUUUGA. For UBC13, GCACAGUUCUGCUAUCGAU, GAGCAUGGACUAGGCUAUA, CAGAUGAUCCAUUAGCAAA, and GCGGAGCAGUGGAAGACCA. For Mcl1, CGAAGGAAGUAUCGAAUUU, AGAACGAAUUGAUGUGUAA, GGACCAACUACAAAUUAAU, GCUACGUAGUUCGGGCAAA.. Cells were transfected with gene-specific siRNAs using the Lonza 4D-Nucleofector X Unit and SF Cell Line 4D-Nucleofector X Kit L (Catalog #: V4XC-2012) according to the manufacturer’s instructions. Knockdown efficiency was verified by immunoblotting 72 h post-transfection. GFP-expressing control cells (shGFP) were used as negative controls for knockdown.

### Immunofluorescence

For immunofluorescence (IF), U266B1^WT^ or U266B1 p62^KD^ cells were treated with 5 or 10 µM AUTAC for either 24 or 48 hours, or with 10 nM CFZ, 5 µM AUTAC, or a combination of AUTAC and CFZ for 16 hours. The treated cells were seeded onto 12-mm-diameter glass coverslips placed in 12-well culture dishes pre-coated with 0.01% poly-L-lysine (Sigma-Aldrich, P4832) for 1 hour. Cells were washed with PBS before fixation in 4% paraformaldehyde for 30 minutes at room temperature (RT). Following fixation, cells were rinsed with PBS and blocked for 1 hour in a blocking buffer containing 5% BSA and 0.1% Triton X-100 in PBS. The cells were then incubated overnight at 4°C with the designated primary antibodies including LAMP1 (Cat. No. 15665S, Cell Signaling Technology), LC3B (Cat. No. 83506S, Cell Signaling Technology), K63 (Cat. No. 14-6077-82 Thermo Fisher), p62/SQSTM1 (Cat. No. 88588 Cell Signaling Technology), and Mcl1 (Cat. No. 94296S, Cell Signaling Technology). After three washes with PBS, the cells were incubated with the corresponding secondary antibodies (Alexa Fluor 488 anti-mouse; Lot: 160324, Jackson ImmunoResearch, Alexa Fluor 555 goat anti-mouse IgG; Lot: WA316324, Invitrogen, and Alexa Fluor 594 goat anti-rabbit IgG; Lot: 1608397, Life Technologies). The secondary antibodies were diluted in antibody solution (5% BSA and 0.1% Triton X-100 in PBS) and incubated for 2 hours at room temperature (RT). Nuclear staining was performed by incubating the cells with 4’,6-diamidino-2-phenylindole (DAPI; Lot: WE3286201, Thermo Scientific) for 3 minutes at RT in the dark. Cells were washed three times with PBS, mounted onto glass slides using a mounting medium (Vectashield), and imaged using a Zeiss Laser Scanning Microsope (LSM) 700 fitted with a 63x oil-immersion lens. Images were analyzed, processed and quantified using ImageJ software. Colocalization was assessed using JACoP with Pearson’s correlation coefficient and Manders’ coefficients (M1, M2). Background subtraction and thresholding were applied consistently across images. The figures were generated using GraphPad Prism software. In all pharmacologic and genetic manipulation experiments, appropriate matched controls were included. DMSO served as the vehicle control for compound treatments. GFP-expressing control cells (sh-GFP) were used as negative controls for knockdown.

### Molecular docking studies

#### Protein preparation.^108^

Crystal structures of human MCL1 (PDB ID: 6YBL), BCL2 (PDB ID: 2XA0), and BCL-XL (PDB ID: 1R2D) were obtained from the Protein Data Bank. Bond orders were assigned, missing hydrogens were added, and protonation states were optimized using Epik at physiological pH (7.0 ± 0.5).^109^ Hydrogen-bonding networks were refined, and crystallographic water molecules were removed except where structurally conserved or involved in key interactions. The prepared protein structures were subjected to restrained energy minimization using the OPLS4 force field^110^, converging heavy atoms to an RMSD of 0.3 Å.

#### Ligand preparation

The AUTAC molecule and the corresponding warhead fragment were prepared using LigPrep with the OPLS4 force field. Ionization states and tautomers were generated using Epik at physiological pH, and all salts were removed. Warhead-only reference structures were prepared using the same protocol to ensure consistent atom typing and charge assignment for subsequent core-constrained docking of the full AUTAC.

#### Docking of warhead and AUTAC

Docking was performed using the Schrödinger’s Glide module in standard precision (SP) mode. For initial warhead docking, receptor grids were generated based on the prepared protein structures. In the case of MCL1, the grid was centered on the co-crystallized selective inhibitor S64315 to define the BH3-binding groove. Identical grid-generation parameters were subsequently applied to BCL2 and BCL-XL to ensure consistent treatment across targets. Docking of the isolated warhead was performed with 100 independent docking runs followed by post-docking minimization. The highest-ranked pose for each protein, based on GlideScore and consistency with canonical BH3-groove interactions, was selected as the reference warhead binding mode. Docking of the full AUTAC molecule was then carried out using a core-constrained protocol to preserve the validated warhead orientation. Atoms defining the rigid warhead pharmacophore were manually selected and constrained to overlay the reference warhead pose with an RMSD tolerance of 0.1 Å. Receptor grids were generated from the warhead-bound protein complexes, centered on the bound warhead and using identical grid dimensions and parameters for MCL1, BCL2, and BCL-XL. Enhanced conformational sampling was employed to allow flexibility of the linker and FBnG degradation tag, and fifty AUTAC poses were generated per target. All poses were subjected to post-docking minimization, ranked by GlideScore, and clustered to identify dominant binding families.

### MM-GBSA binding energy calculations

Binding free energies of AUTAC–protein complexes were estimated using the Molecular Mechanics/Generalized Born Surface Area (MM-GBSA) method. MM-GBSA calculations were performed using the **OPLS4 force field** in combination with the **VSGB implicit solvent model**. Prior to energy evaluation, each protein–ligand complex was subjected to restrained energy minimization to relieve local strain while preserving the docked geometry. Binding free energies (ΔG_bind) were computed as the difference between the minimized complex energy and the sum of the minimized energies of the isolated protein and ligand. The total binding free energy was decomposed into **van der Waals (**Δ**G_vdW)**, and **electrostatic (**Δ**G_electrostatic)** contributions. Average MM-GBSA values were reported as the mean of the top five poses for each target.

### Solvent-accessible surface area (SASA) analysis of FBnG exposure

Molecular surfaces were generated using a **probe radius of 1.4 Å**, corresponding to a water molecule. For each target, the FBnG tag was explicitly selected, and its SASA was calculated in both the **isolated (free) state** and the **protein-bound complex** using identical surface generation parameters. FBnG exposure in the bound state was quantified as the absolute SASA and as a **fraction of the free-state SASA**. Differences in FBnG accessibility upon binding (ΔFBnG) were calculated as FBnG_free − FBnG_bound.

### Statistical analysis and reporting

Unless otherwise indicated, data are presented as mean ± SEM from at least three independent biological replicates, each performed with technical triplicates (or six technical replicates for cell viability assays). Statistical analyses were performed using GraphPad Prism 10. Statistical significance of each condition compared to the indicated control or treatment was determined using unpaired, two-sided Student’s t-tests or two-way ANOVA with Tukey’s or Sidak’s post hoc tests, as appropriate. A P value < 0.05 was considered statistically significant. Statistical significance levels are indicated in the figures as follows: *P < 0.05; **P < 0.01; ***P < 0.001; and ****P < 0.0001.

## Data and Code Availability

No high-throughput sequencing or omics datasets requiring deposition were generated. Source data underlying figures are available from the corresponding author upon reasonable request.

No custom code was generated. Molecular docking was performed using Schrödinger software (Glide/LigPrep/Epik/MM-GBSA; OPLS4). Image analyses were performed in ImageJ; statistics and plots were generated in GraphPad Prism 10.

## ACKNOWLEDGEMENTS

This work was supported by a NIH/NIGMS grant R01GM132396 (S.K.R). We thank Gugan Thuduppathy for helpful discussions. We thank Bin Hu, and Wang Li for their assistance with the mouse studies. Services and products in support of the research project were generated by the Virginia Commonwealth University Cancer Mouse Models Core Laboratory and Microscopy Shared Resource, supported, in part, with funding to the Massey Cancer Center from NIH-NCI Cancer Center Support Grant P30 CA016059.

## CONFLICT OF INTEREST STATEMENT

The authors declare no conflict of interest.

## DATA AVAILABILITY STATEMENT

All data that support the findings of this study are available from the corresponding author upon reasonable request.

## FIGURE LEGENDS

**Supplementary Figure 1. Structural comparison and docking of AUTAC warhead in Mcl1, Bcl2, and Bclxl. (A)** Sequence alignment of human Mcl1, Bcl2 and Bclxl; **(B)** Structural overlay of Mcl1 vs Bcl2 vs Bclxl BH3- binding grooves. Mcl1 (PDB ID: 2PQK), Bcl2 (PDB ID: 2XA0), Bclxl (PDB ID 1R2D) and BH3-peptides are shown as grey, yellow, orange and salmon cartoons respectively. **(C)** Highest-ranked docking pose of the warhead in (**C1**) Mcl1, (**C2**) Bcl2, and (**C3**) Bclxl. Proteins are shown as cartoons (Mcl1 grey, Bcl2 yellow, Bclxl orange). The warhead is shown as salmon sticks. Key pocket residues are shown as sticks, and hydrogen-bonding interactions are indicated by dashed lines. **(D)** Representative florescence microscopy images and their quantification for U266B1 cells subjected to starvation for 24h, then the cells stained with DAPI and lysotracker stain. All images were generated under the same magnification (20X). **(E)** Western blot analysis for LC3I/II protein in U266B1 starved and treated with CQ (20 μM) for 24h. GAPDH was used as the loading control.

Three biological replicates for each cell line were used to perform Western blotting. The uncropped blots are shown in the Supplemental Materials.

**Supplementary Figure 2. AUTAC promotes lysosomal and autophagic association of Mcl1 in U266B1 cells. (A)** Western blot analysis for mTOR, p53, HSP70, HSP60, ERK, and ULK1 proteins in U266B1 treated with AUTAC 5 μM for 24h. GAPDH was used as the loading control. **(B)** Confocal images with their quantification for Mcl1, LAMP1 and LC3B proteins in U266B1 cells treated with AUTAC (5 μM) for 24 and 48h (scale 10 μM).

Three biological replicates for each cell line were used to perform Western blotting. The uncropped blots are shown in the Supplemental Materials. Statistical significance of each condition compared to the indicated control or treatment was determined using unpaired Student’s t-test or two-way ANOVA with Tukey or Sidak post hoc tests, as appropriate. Data are represented as mean ± SEM. Significance levels are indicated as follows: * p < 0.05, ** p < 0.01, *** p < 0.001 and **** p < 0.0001.

**Supplementary Figure 3. AUTAC selectively degrades Mcl1 and subsequently induces autophagy through a Beclin1–dependent axis.(A)** Western blot analysis for Mcl1, and LC3I/II proteins in U266B1 treated with Mcl1 AUTAC (5μM) or MetAP2 AUTAC (10μM) for 24h. GAPDH was used as the loading control. **(B)** Western blot analysis for LC3I/II protein in U266B1 treated with CQ (20 μM) and/or Mcl1 AUTAC (5μM) or MetAP2 AUTAC (10μM) for 24h. GAPDH was used as the loading control. **(C)** Western blot analysis for Mcl1, mTOR, p-mTOR, T-S6, and p-S6 proteins in U266B1 treated with Mcl1 AUTAC (5μM) and rapamycin (100nM) for 24h. GAPDH was used as the loading control. **(D)** Western blot analysis for LC3I/II protein in U266B1 treated with CQ (20 μM) and/or Mcl1 AUTAC (5μM), or rapamycin (100nM) for 24h. GAPDH was used as the loading control. **(E)** Western blot analysis for LC3I/II protein in U266B1 treated with CQ (20 μM) and/or Mcl1 moiety (10μM), or FBNG (10μM) for 24h. GAPDH was used as the loading control. **(F)** Western blot analysis for Mcl1, and LC3I/II proteins in U266B1^WT^, and U266B1 Mcl1 ^KD^ treated with CQ (20 μM) and/or AUTAC (5μM) for 24h. GAPDH was used as the loading control. **(G)** Western blot analysis for LC3I/II protein in U266B1 treated with CQ (20 μM), AUTAC (5 μM), and N-acetyl cysteine (NAC) (0.5 μM) for 24h. GAPDH was used as the loading control. **(H)** Western blot analysis for Beclin1, and p-Beclin1 (Ser15) proteins in U266B1 treated with AUTAC (5 μM) for 24h. GAPDH was used as the loading control.

Three biological replicates for each cell line were used to perform Western blotting. The uncropped blots are shown in the Supplemental Materials.

**Supplementary Figure 4. Autophagy induction via starvation augments the Mcl1 degradation ability of AUTAC. (A)** Western blot analysis for Mcl1 protein in U266B1 treated with AUTAC (5 μM), NSC697923 (1μM) and/or C25-140 (10 and 30 μM) for 24h. GAPDH was used as the loading control. **(B)** Western blot analysis for p62, Mcl1, and LC3I/II proteins in U266B1 starved and treated with AUTAC (5 μM) for 24h. GAPDH was used as the loading control.

Three biological replicates for each cell line were used to perform Western blotting. The uncropped blots are shown in the Supplemental Materials.

**Supplementary Figure 5. Carfilzomib induces a cytoprotective autophagy in multiple myeloma cells. (A)** Representative florescence microscopy images with their quantification for U266B1 cells treated with carfilzomib (10 nM) for 24h, then the cells stained with DAPI and lysotracker stain. All images were generated under the same magnification (20X). **(B)** Western blot analysis for LC3I/II protein in MM.1S and RPMI-8226 cells treated with carfilzomib (10 nM) and/or CQ (20 μM) for 16h. GAPDH was used as the loading control. **(C)** Western blot analysis for LC3I/II protein in U266B1 and U266B1^R^ treated with carfilzomib (10 nM) and/or CQ (20 μM) or bafilomycin (10 nM) for 24h. GAPDH was used as the loading control. **(D and E)** The percentage of cellular viability for **(D)** U266B1 and **(E)** U266B1^R^ cells treated with CQ (20 μM) or bafilomycin (10 nM) for 72h, then the indicated concentrations of CFZ were added for the last 24h. The synergy between CQ, or BAF and CFZ were determined using Bliss score.

Three biological replicates for each cell line were used to perform cell viability assay, and Western blotting. In each independent replicate of the synergy assay, a single well was used for each drug combination. The uncropped blots are shown in the Supplemental Materials. ns indicates non-statistical significance, while **** p < 0.0001 indicates statistical significance of each condition compared to the indicated control/treatment as determined using two-way ANOVA with Sidak’s post hoc test.

**Supplementary Figure 6. Mcl1 contributes to proteasome inhibitor resistance in multiple myeloma cells. (A)** Viability of U266B1^WT^, U266B1^R^, and U266B1 Mcl1 overexpression (OE) cells treated with the indicated concentrations of CFZ for 24h. **(B)** Western blot analysis for Mcl1 protein in U266B1^WT^, U266B1^R^, and U266B1 Mcl1 ^OE^. GAPDH was used as the loading control. **(C)** Western blot analysis for Mcl1, LC3I/II, cleaved PARP, and cleaved caspase 3 proteins in U266B1^WT^ and U266B1 Mcl1^OE^ cells treated with AUTAC (5 μM) for 48h, then CFZ (10 nM) was added for the last 24h. GAPDH was used as the loading control. **(D)** The percentage of cellular viability for U266B1^WT^ and U266B1 Mcl1^OE^ cells treated with AUTAC (5 μM) for 72h, then the indicated concentrations of CFZ were added for the last 24h.

Three biological replicates for each cell line were used to perform cell viability assay, and Western blotting. The uncropped blots are shown in the Supplemental Materials.

## Notes

### Competing Interest Statement

The authors have declared no competing interest.

### Summary of Updates

More mechanistic details included. Also in vivo xenograft studies included.

