## Supplementary figures and images for "Proteasome Inhibition Enhances Lysosome-mediated Targeted Protein Degradation"

### Supp Fig 1

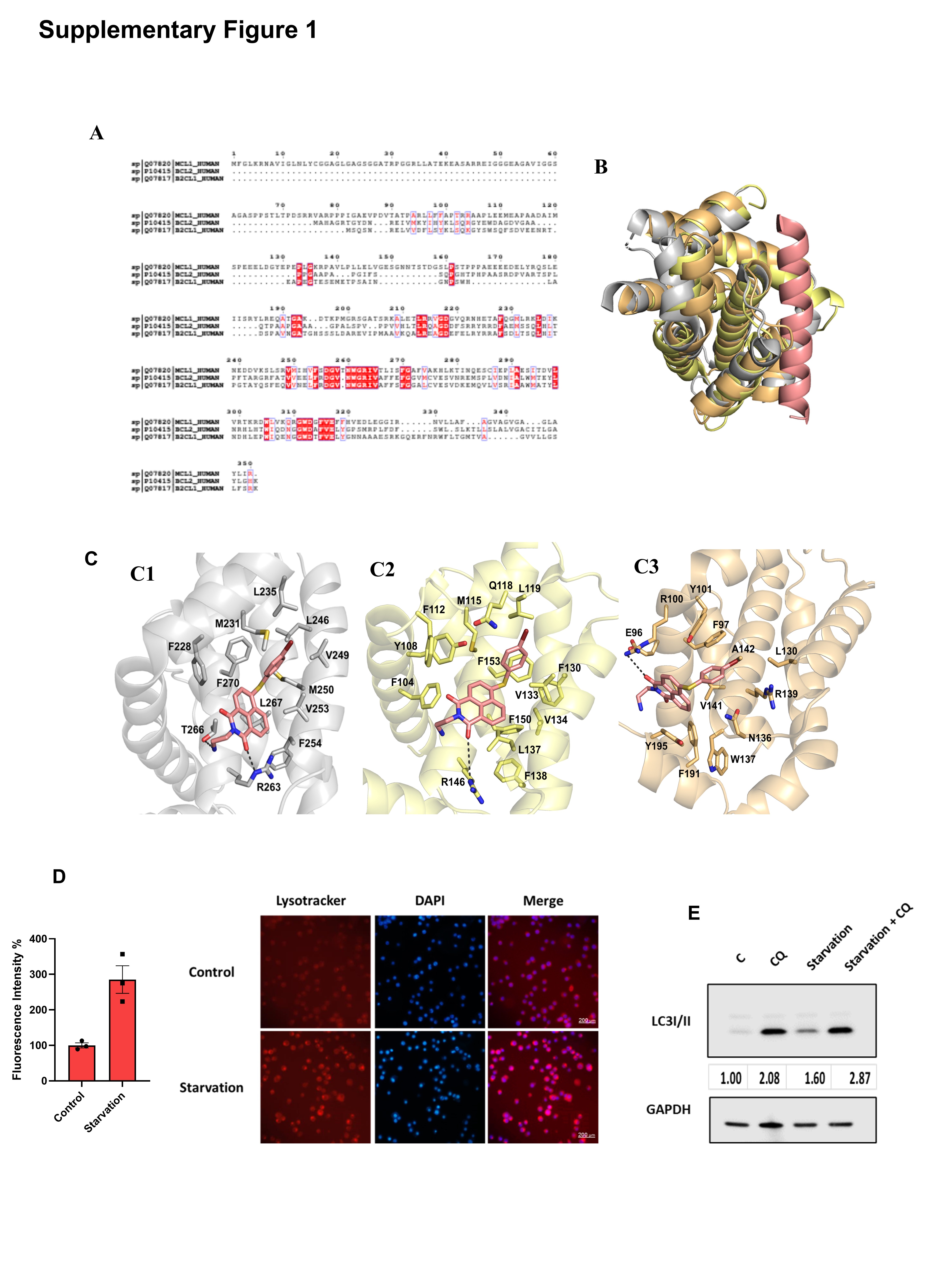

### Supp Fig 2

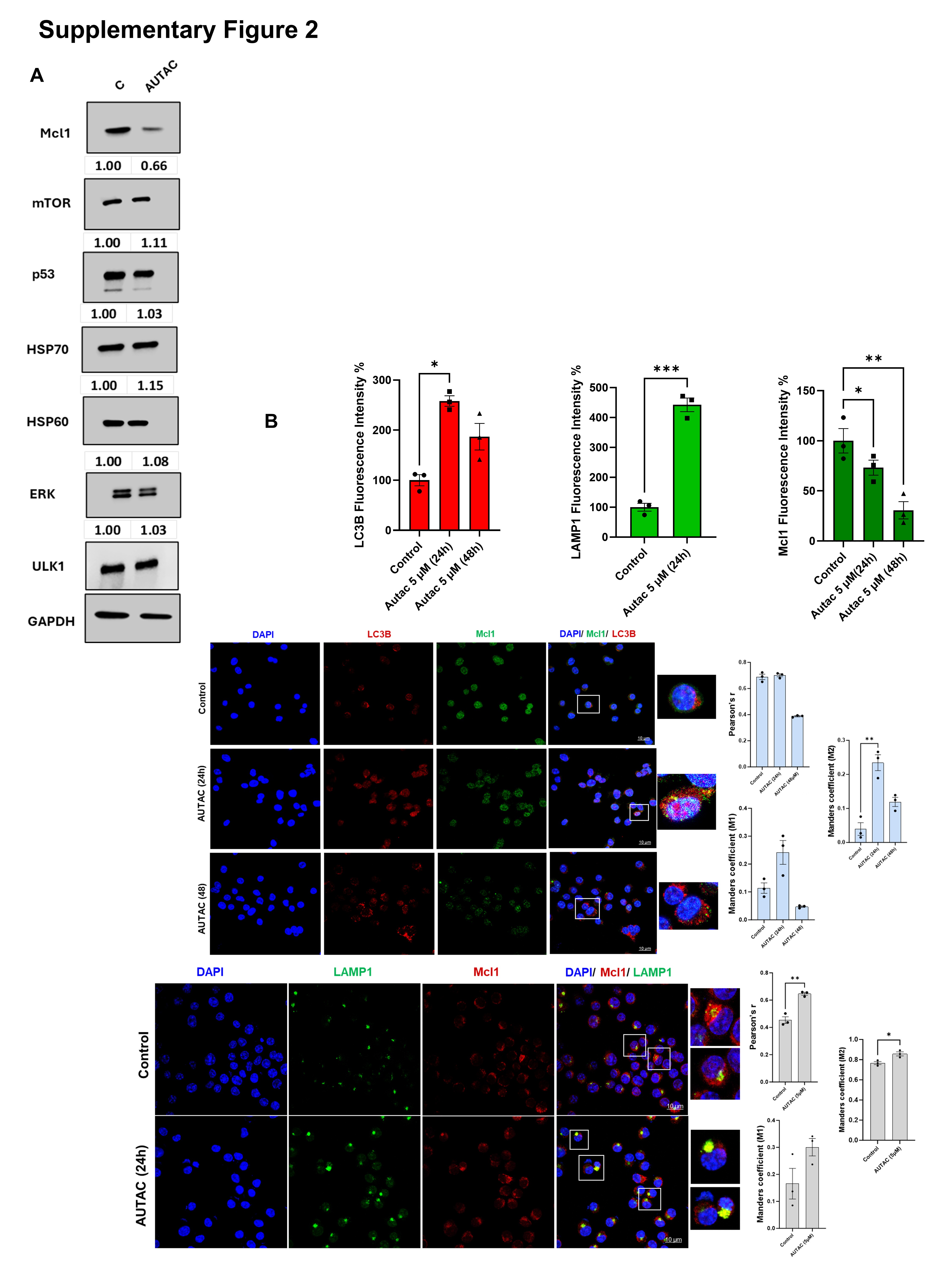

### Supp Fig 3

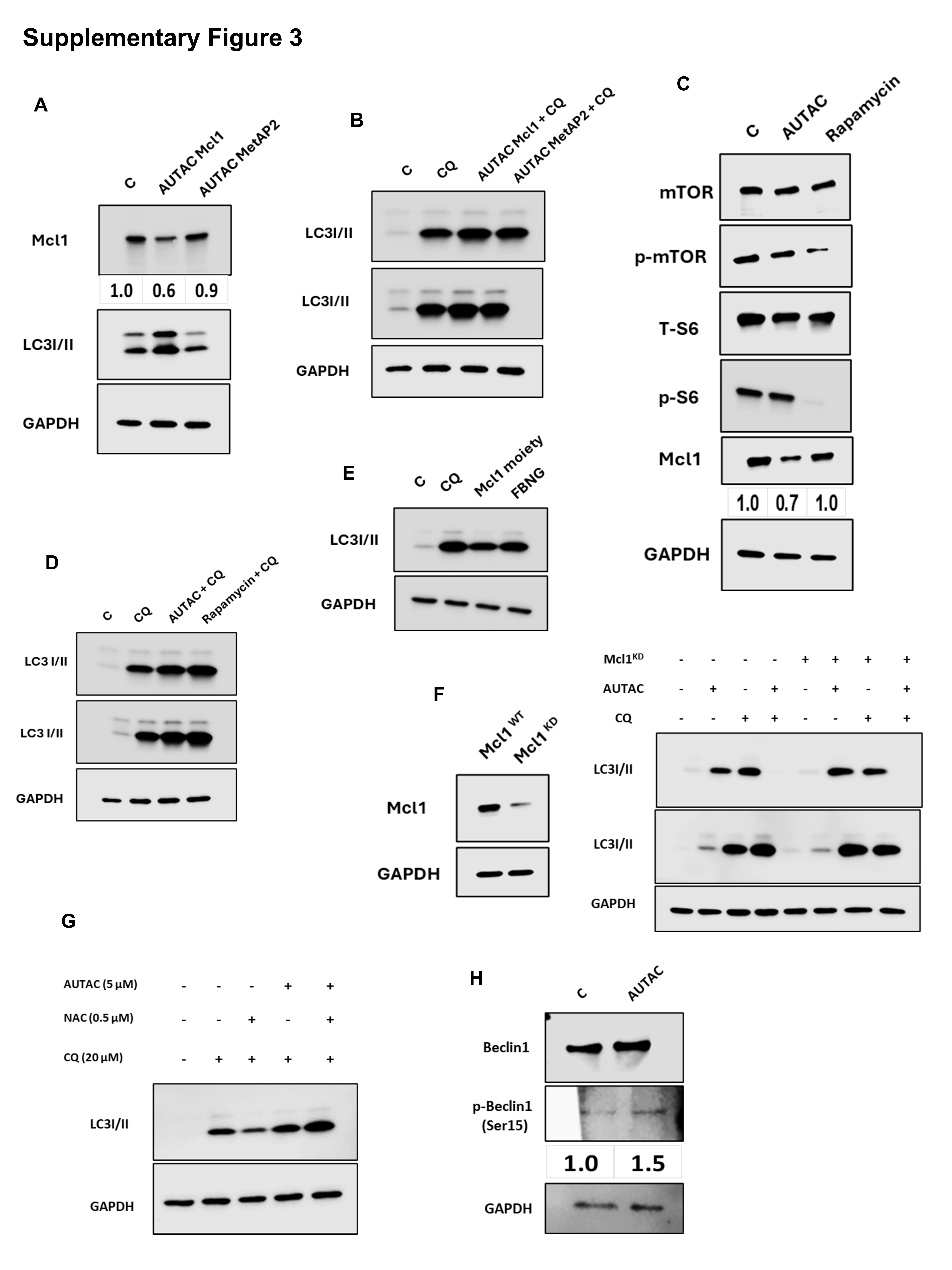

### Supp Fig 4

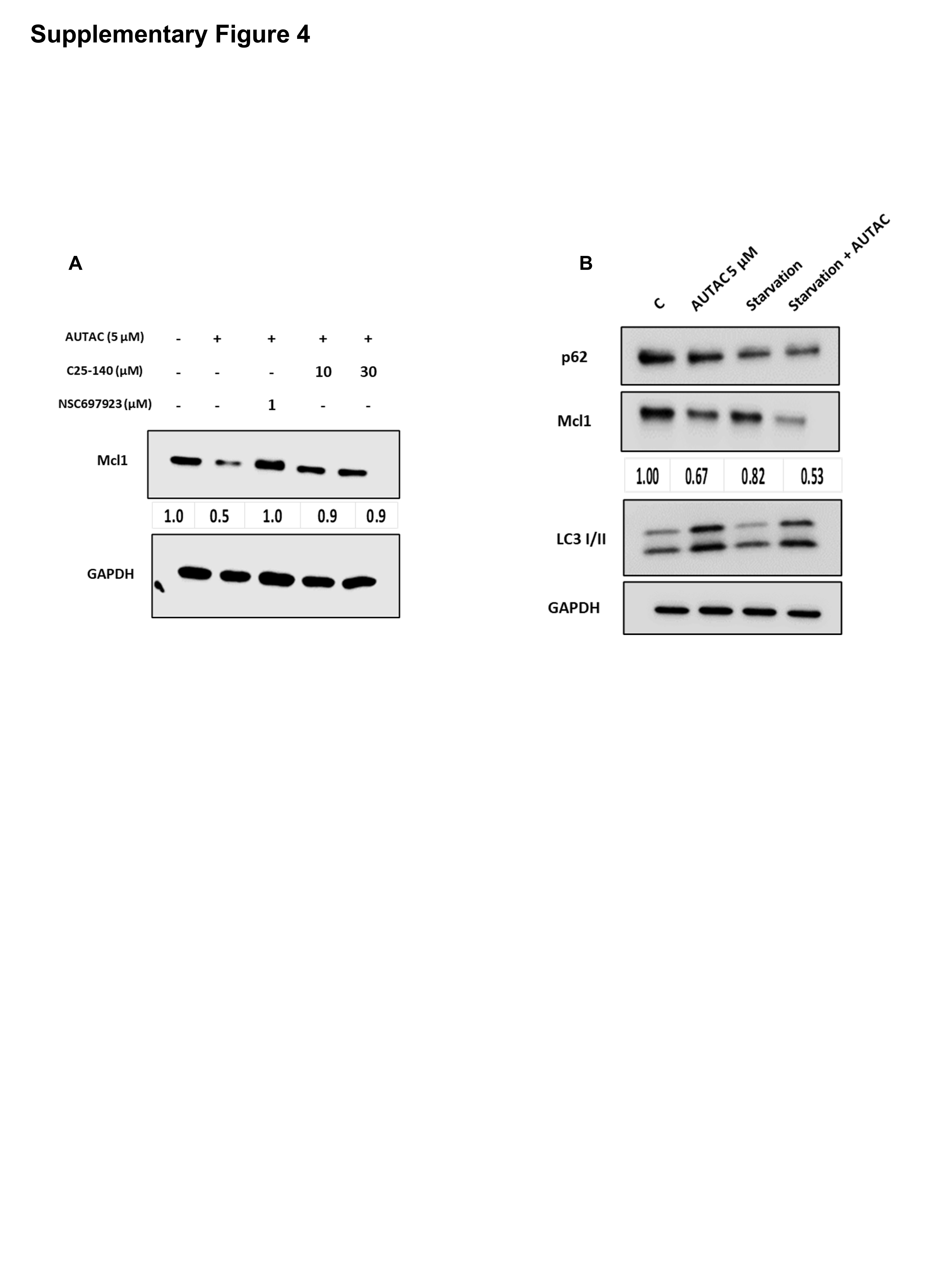

### Supp Fig 5

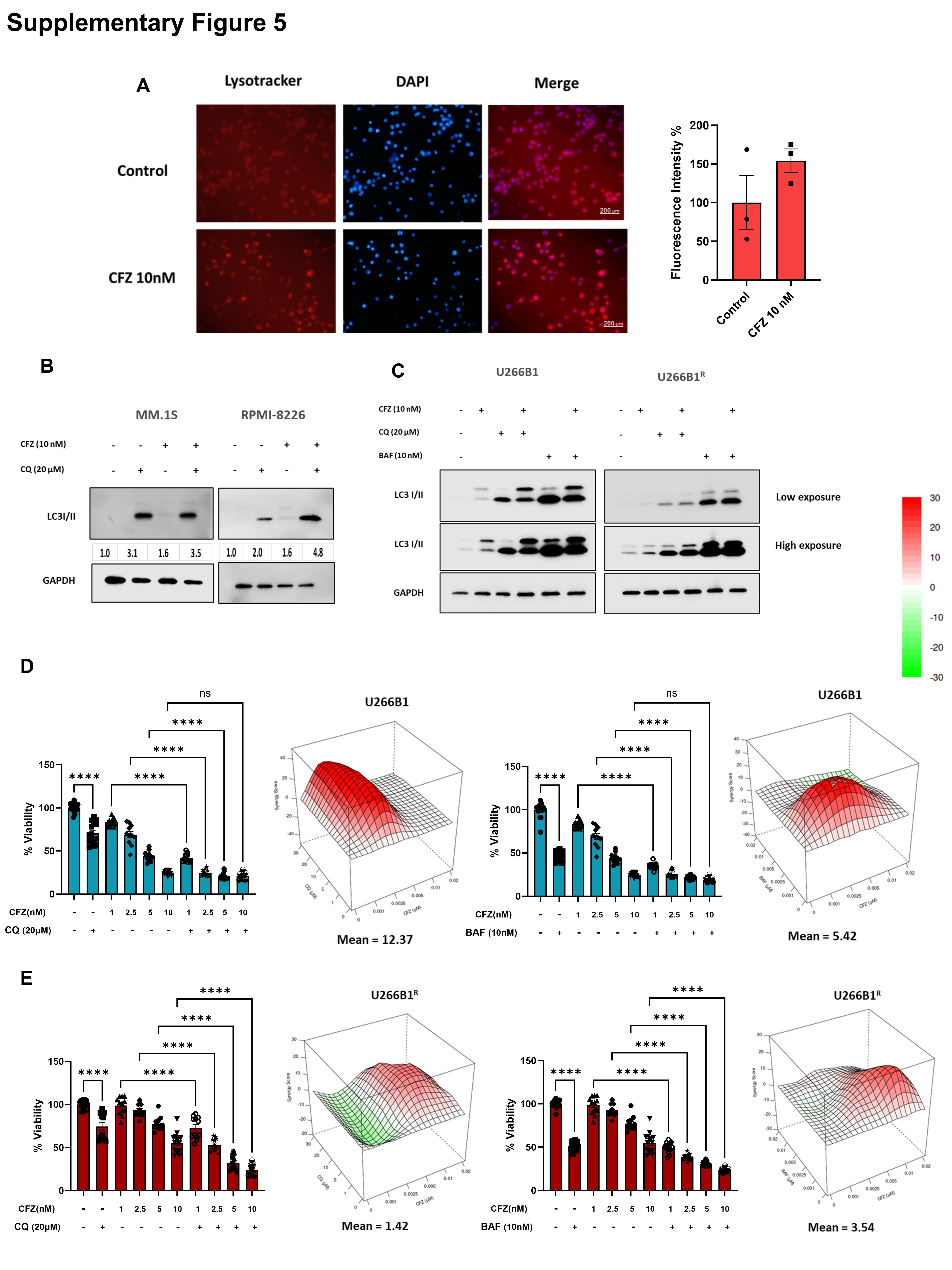

### Supp Fig 6

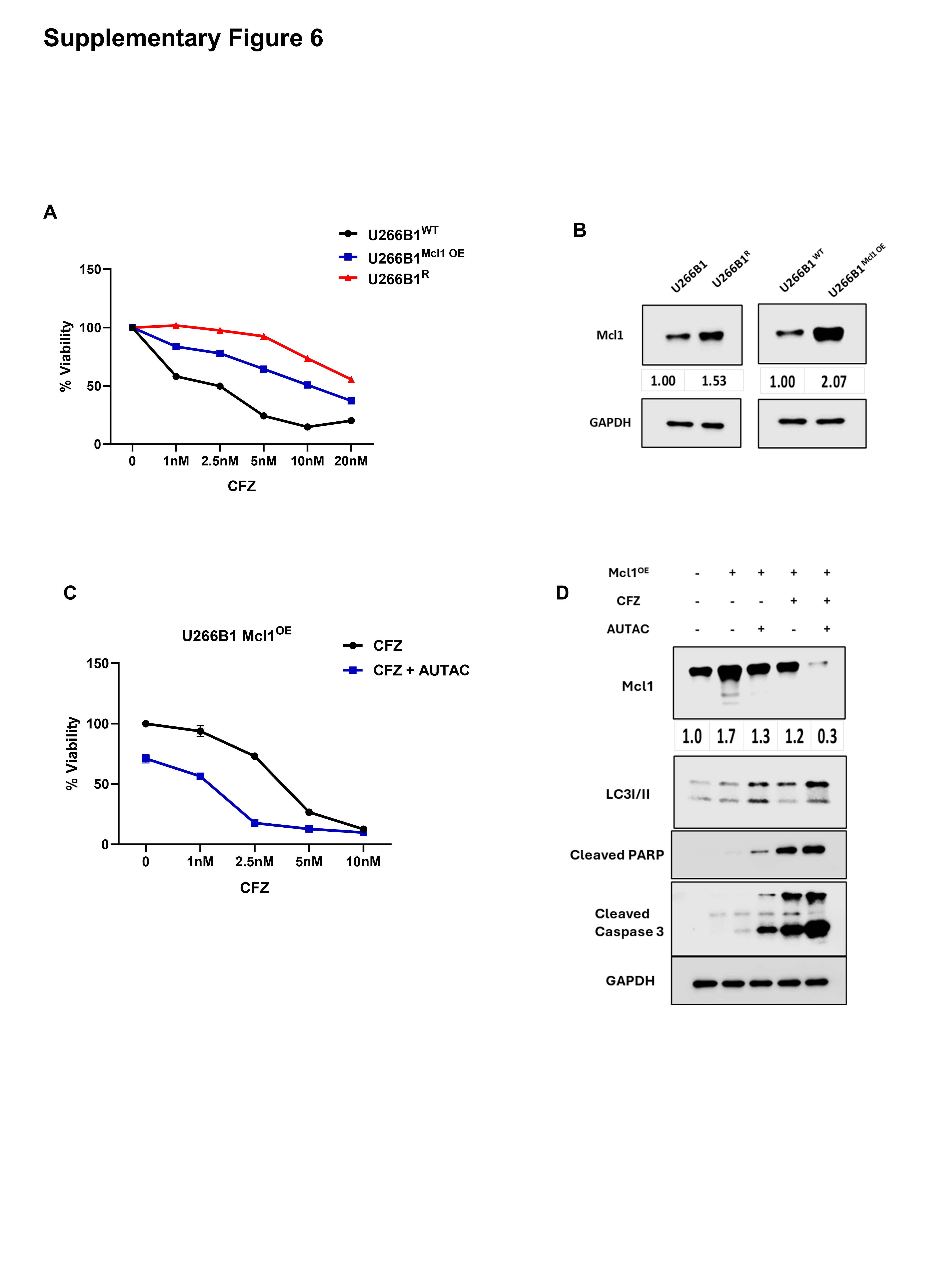
